# Gangliosides Modulate the Secretion of Extracellular Vesicles and Their Misfolded Protein Cargo

**DOI:** 10.1101/2025.04.03.647108

**Authors:** John Monyror, Vaibhavi Kadam, Luis Carlos Morales, Diego Ordóñez, Jeremies Ibanga, Asifa K. Zaidi, Emily McNamara, Desmond Pink, Magnus Stenlund, Ken Reyes, Aislinn D. Maguire, Jing Huang, Leonardo M. Cortez, Valerie Sim, Sue-Ann Mok, Elena Posse de Chaves, Simonetta Sipione

## Abstract

Gangliosides are glycosphingolipids that play an integral role in cell signaling and provide neuroprotection. While present on extracellular vesicles (EVs) – key mediators of intercellular communication – their role in EV biogenesis remains unclear. Here, we identify gangliosides, both endogenously synthesized and exogenously administered, as key modulators of EV biogenesis, with the specific composition of their glycan headgroup and the presence or absence of sialic acid and N-acetyl-D-galactosamine residues dictating whether they promote or inhibit EV biogenesis. We show that GM1 and other complex gangliosides enhance EV secretion, while disruption of ganglioside synthesis impairs it. GM1 supplementation restores EV secretion in Huntington’s disease (HD) fibroblasts and HD cell models that have lower than normal levels of gangliosides, and in cells with a genetic block of ganglioside synthesis that models rare early-onset neurodegenerative diseases. Notably, GM1 also enhances EV-mediated secretion of pathogenic misfolded proteins, including mutant huntingtin (mHTT), α-synuclein and tau, reducing intracellular burden and providing mechanistic insight into the mHTT-lowering effects of GM1 treatments in HD models. Our findings shed light on the neuroprotective roles of gangliosides and highlight their potential for therapeutic exploitation in misfolded protein disorders.

## Introduction

Extracellular vesicles (EVs) are membrane-bound particles ranging from 30-1000 nm in diameter, released by cells into the extracellular space^1^. They originate either from intraluminal vesicles of multivesicular bodies (MVBs) when these fuse with the plasma membrane (exosomes), or from direct budding of the plasma membrane (ectosomes). EVs carry a large repertoire of macromolecules involved in their biogenesis and functions, enabling them to play critical roles in intercellular communication and the transfer of macromolecules and organelles between cells, under both physiological and pathological conditions^1–4^. In the brain, EV-mediated molecular transfer and signaling influence development, synaptic plasticity, proteostasis and microglia activity^5–7^.

EVs also play a central role in misfolded protein diseases such as Alzheimer’s disease (AD), Parkinson’s disease (PD), and Huntington’s disease (HD). The release of EVs carrying pathogenic misfolded proteins helps alleviate proteotoxic stress in vulnerable cells and promotes the clearance of toxic proteins^8,9^. However, EVs can also contribute to the intercellular transfer of misfolded proteins between neurons, promoting their prion-like spread throughout the brain^10–13^. A central role of EVs in neurodegenerative diseases was further suggested by a large systems biology study that identified the EV pathway and protein network as among the most highly affected in multiple misfolded protein diseases^14^. These findings raise critical questions about how misfolded protein diseases impact EV biogenesis and function, and how these alterations may, in turn, influence the clearance and propagation of pathogenic proteins.

Neurodegenerative misfolded protein diseases are associated with significant alterations in lipid metabolism^15^, which contribute to both neurodegeneration and neuroinflammation^16–18^. Many of the affected lipids play key roles in EV biogenesis^19^, including ceramide, cholesterol, phosphatidylinositol, phosphatidylethanolamine and phosphatidylserine^20–22^. Notably, several proteins involved in EV biogenesis localize to membrane microdomains enriched with cholesterol, sphingolipids and gangliosides^23–26^. These microdomains serve as platforms where lipid composition influences protein binding, clustering and activity, ultimately impacting EV biogenesis^26–28^. Many of these lipids and proteins are incorporated and enriched into EVs themselves^1^. Thus, lipid abnormalities in neurodegenerative diseases may disrupt EV biogenesis and functions, impairing EV-mediated cell-cell communication and proteostasis.

Gangliosides - sialic acid-containing glycosphingolipids highly abundant in the central nervous system^29,30^ - are another key component of EV membranes^31^. They localize to membrane microdomains rich in EV-associated lipids and proteins^32^, and exhibit notable biophysical properties, including an amphiphilic nature, asymmetric distribution within membrane leaflets, and the ability to induce positive curvature in cell membranes – a feature shared by other EV-promoting lipids^33–35^. These characteristics suggest a role for gangliosides in EV formation, yet this possibility remains largely unexplored. Addressing this knowledge gap is particularly relevant given that brain ganglioside levels are altered in several common neurodegenerative diseases^29,36–40^. For example, ganglioside GM1 synthesis is decreased in PD and HD, while loss of function mutations in ganglioside biosynthetic enzymes result in the lack of complex gangliosides in a form of hereditary spastic paraplegia and other rare early-onset neurodegenerative disorders^41–44^. Importantly, exogenous GM1 supplementation has been shown to alleviate symptoms^40,45–48^ and reduce brain levels of mutant huntingtin (mHTT)^45^ and alpha-synuclein^49^ in models of HD and PD, respectively. However, whether altered ganglioside levels contribute to impaired EV secretion, and how GM1 supplementation influences this process remains unknown.

Here, we describe a novel role for gangliosides, both endogenously synthesized and exogenously administered, in modulating EV secretion. We demonstrate that gangliosides influence EV release by neuronal and non-neuronal cells in a bidirectional manner, depending on their specific glycan structure. Furthermore, we show that reduced cellular ganglioside levels impair the ability of cells to dispose of mHTT via the EV pathway, whereas GM1 administration enhances EV secretion, facilitating the export of misfolded proteins and decreasing intracellular mHTT burden. These findings provide new insights into the neuroprotective effects of GM1 and uncover a previously unrecognized role for gangliosides in EV biogenesis, with significant implications for the pathogenesis and treatment of neurodegenerative diseases.

## Results

### Administration of ganglioside GM1 promotes the secretion of EVs from murine and human neuronal and non-neuronal cells

To assess the impact of gangliosides on EV secretion, we began by examining the effects of ganglioside GM1 on neuronal cells. GM1 administration results in ganglioside uptake and membrane enrichment in treated cells^40,50^. To facilitate direct analysis of EVs secreted into the culture medium while minimizing sample handling and EV loss, we used a state-of-the-art imaging flow cytometry (IFC) platform optimized and validated for the analysis of fluorescent EVs^51,52^ produced by cells pre-labelled with a lipophilic dye (DiD or DiI, as indicated), which is incorporated in all cell membranes, including those of EVs^53^. Apoptotic bodies and cell debris were removed from the conditioned medium by centrifugation at 2,000 x g (cleared conditioned medium, CCM) before EV analysis^53^. In our initial experiments, neuroblastoma N2a cells were incubated with 50 μM GM1 for 6 h, followed by washing and EV collection in GM1-free culture medium for 16 h. In these conditions, cells treated with GM1 secreted 50% more EVs compared to untreated controls (Fig. 1a and Supplementary Fig. 1). Notably, the increase in EV secretion was significantly higher when GM1 remained present throughout the 22h EV collection period (Fig. 1b). This effect is likely due to the continuous stimulatory action of GM1 on EV secretion, compared to conditions where the ganglioside was removed, and its cellular levels were allowed to return to baseline due to ganglioside catabolism and cell division. Similar results were observed when EVs were isolated from the CCM by size-exclusion chromatography (SEC, Fig. 1c) or ultracentrifugation (Fig. 1d), two widely used methods^54,55^. Western blot analysis of isolated EVs confirmed the expected enrichment for the EV marker Flotillin-1 (Fig 1d).

**Figure 1.**
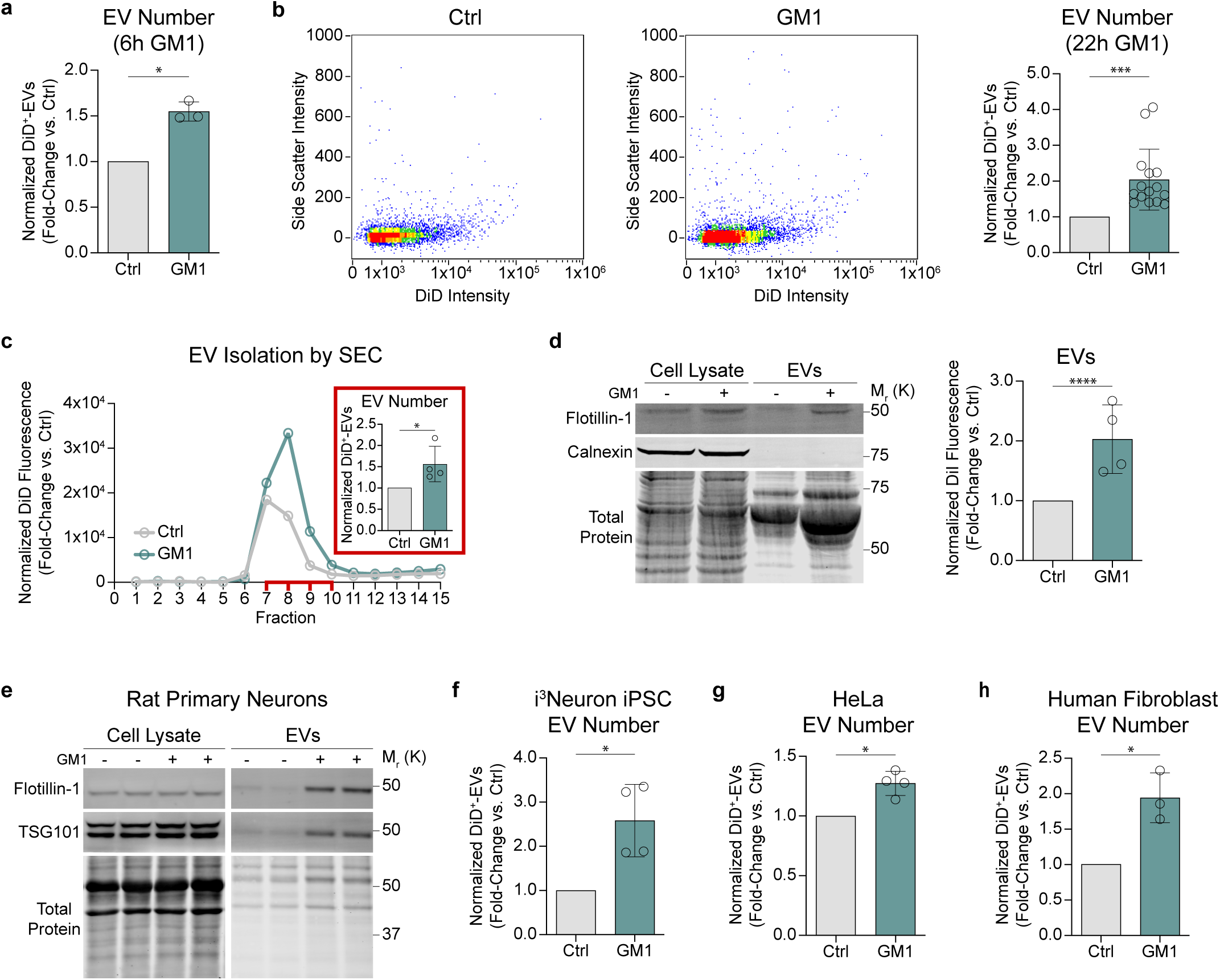
Administration of ganglioside GM1 promotes the secretion of EVs from murine and human cells of different origins. **a**. N2a cells were treated with 50 µM GM1 for 6 h. GM1 was washed-off the cells and EVs were collected in serum-free medium supplemented with N-2 for 16 h and counted by IFC (n=3 independent experiments). **b.** Representative IFC density dot plots and quantification of DiD^+^-EVs secreted by control or GM1-treated N2a cells. Cells were treated with 50 µM GM1 throughout a 22 h EV collection period (n=15 independent experiments). **c.** Representative size-exclusion chromatogram and quantification (inset) of EVs secreted by N2a cells treated with GM1 or vehicle for 6 h. DiD fluorescence in each fraction was normalized to cellular protein content and the peaks correspond to EV-rich fractions (7-10). The inset shows the number of DiD^+^-EVs in the EV-rich fractions measured by IFC and normalized to total cellular protein (n=4 independent experiments). **d.** N2a cells were treated with 50 µM GM1 for 18 h. GM1 was washed off the cells and EVs collected in the medium for the subsequent 24 h were isolated by ultracentrifugation. (Left) Representative immunoblot of the EV marker flotillin-1 and calnexin to confirm absence of cell contaminants. (Right) DiI fluorescence in the EV pellet was measured by fluorometry and normalized to total cellular protein content (n=4 independent experiments). **e.** DIV 15 embryonic rat cortical neurons were treated with 50 µM GM1 in growth medium for 72 h. Equal fractions of ultracentrifugation-isolated EV pellets derived from control and GM1-treated neurons were immunoblotted (n= two independent primary neuronal cultures). **f.** Human i^3^Neurons were treated with 50 µM GM1 throughout a 22 h EV collection period (n=4 independent experiments). **g.** HeLa cells were incubated with 50 µM GM1 for 18 h. GM1 was washed off the cells and EVs were collected for 24 h (n=4 independent experiments). **h.** Human fibroblasts from a healthy subject were incubated with 50 µM GM1 for 6 h. GM1 was washed-off the cells and EVs were collected for 16 h (n=3 independent experiments). (**b, e-h**) The number of DiD^+^-EVs in the CCM was measured by IFC and normalized to total cellular protein content. Bars represent mean ± SD. \**p*<0.05, \*\*\**p*<0.001 by two-tailed paired *t*-test.

We then extended our investigations to determine whether the effects of GM1 on EV secretion are cell type- and species-dependent. GM1 treatment significantly increased EV secretion in primary rat neurons and human induced pluripotent stem cell (iPSC)-derived i^3^Neurons, as measured by EV isolation by ultracentrifugation and immunoblotting for the EV markers Flotilin-1 and TSG-101 (for rat neurons, Fig. 1e), and by IFC analysis of fluorescent EVs in the medium of DiD-labelled human i^3^Neurons (Fig. 1f). The stimulatory effects of GM1 on EV secretion also extended to non-neuronal cells, including HeLa cells (Fig. 1g) and primary human fibroblasts (Fig. 1h). Altogether, through complementary approaches for EV isolation and analysis, our data demonstrate that cell enrichment with GM1 enhances EV secretion in a wide range of cells, highlighting a novel role for ganglioside GM1 in EV biogenesis.

### GM1 does not affect the size and tetraspanin profile of EVs, but increases their GM1 content

To gain insights into the mechanism of action of GM1 and determine whether cell enrichment with GM1 affects a specific EV subpopulation -e.g. small versus large EVs with presumably different subcellular origin, or EV subpopulations characterized by specific tetraspanin profiles^56,57^ - we analyzed EV size by nanoparticle tracking analysis (NTA)^58^, dynamic light scattering (DLS)^59^ and ExoView^60^, and profiled EV tetraspanins using ExoView microchips^60,61^ (Fig. 2b). NTA analysis of EVs in the CCM from GM1-treated and untreated N2a cells showed that GM1 treatment did not alter EV size distribution (Fig. 2a). The EVs detected were mostly small EVs with a diameter of around 150 nm, which is in the size range of both exosomes and ectosomes^1^. We obtained similar results by DLS analysis after fractionating EVs by asymmetric flow field-flow fractionation (Supplementary Fig. 2a). GM1 treatment did not affect the size of EVs secreted by human iPSC-derived i^3^Neurons, human fibroblasts or HeLa cells (Fig. 2c), although, as expected, the average size of EVs measured in these cells by ExoView was smaller, likely due to EV dehydration during ExoView sample preparation^62^.

**Figure 2.**
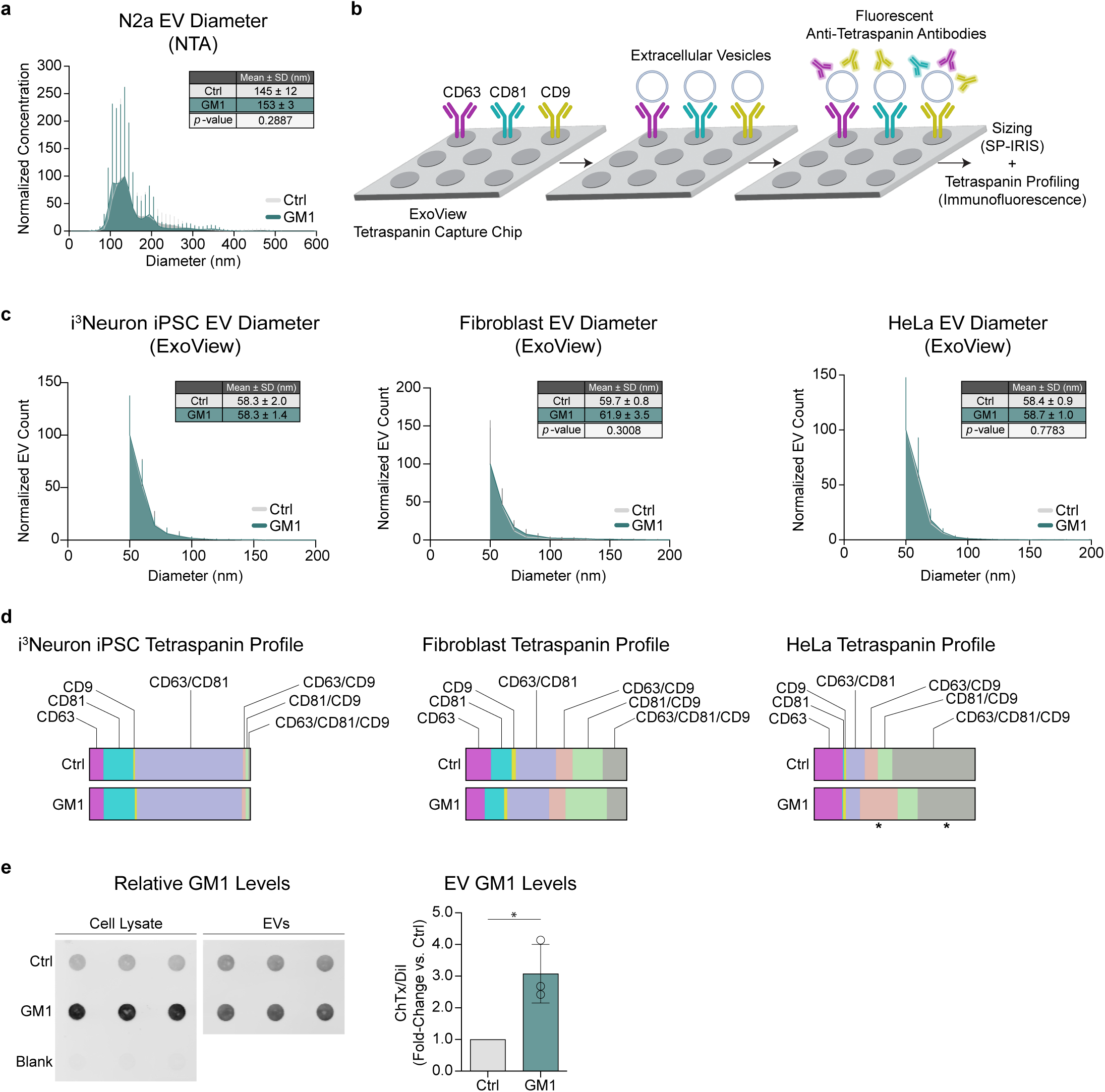
Cell treatment with GM1 preserves EV size and tetraspanin profile but increases EV GM1 content. **a**. Particle size distribution profile of EVs secreted by N2a cells treated with GM1 compared to untreated cells, as measured by NTA. EV concentration was normalized to a common scale. The table reports the mean particle diameter ± SD and *p*-values calculated by two-tailed paired *t*-test (n=3 independent experiments). **b.** Schematic of the ExoView tetraspanin-capture chip and the method used to capture and profile EVs. EVs are captured onto chips by anti-tetraspanin antibodies printed onto the chips. EVs are then labeled with fluorescent anti-tetraspanin antibodies and imaged for tetraspanin profiling and sizing by SP-IRIS. **c.** Particle size distribution profile of EVs secreted by human i^3^Neurons, human fibroblasts and HeLa cells treated with GM1compared to untreated cells, as detected by SP-IRIS. EV counts were normalized to a common scale. The table inserts report the mean particle diameter ± SD and *p*-values obtained by two-tailed paired *t*-test (n=2-3 independent experiments). **d.** Tetraspanin profile of EVs secreted by human i^3^Neurons, human fibroblasts and HeLa cells treated with GM1 compared to untreated cells. Colored bars show the proportion of EVs bearing the indicated tetraspanin combinations. \**p*<0.05 by two-way ANOVA of probit-transformed data with Šídák’s post-hoc test (n=2-3 independent experiments). **e.** Representative dot blot of cholera toxin B binding to quantify GM1 in cell lysates and EVs from N2a cells treated with 50 µM GM1 for 6 h, compared to untreated cells. GM1 was washed off and EVs were collected for 16 h and isolated by SEC. The graph shows the densitometric analysis of cholera toxin B binding to EV fractions, normalized to DiI EV fluorescence. Bars are mean ± SD. \**p*<0.05 by two-tailed paired *t*-test (n=3 independent experiments).

Next, we analyzed the tetraspanin profile of EVs, as this has been proposed to vary depending on the intracellular origin of EVs^63,64^. While both human fibroblasts and i^3^Neurons produced EVs with quite different tetraspanin profiles, the proportion of EVs carrying the major tetraspanins CD63, CD81 and CD9 in different combinations was not significantly altered by GM1 treatment (Fig. 2d). This suggests that GM1 does not promote the secretion of specific EV subpopulations but rather enhances total EV secretion, maintaining the specific EV tetraspanin profile of each cell type. In HeLa cells, however, GM1 treatment increased the proportion of EVs bearing both CD63 and CD9 (from 8.1% in the medium of untreated cells to 23.4% of total EVs in the medium of GM1-treated cells), at the expense of CD63/CD9/CD81 triple positive EVs (from 51.4% to 35.7%) (Fig. 2d). Thus, while GM1 promotes EV secretion in all cells, the specific subpopulation secreted might be cell-type dependent, suggesting GM1 might affect fundamental shared mechanisms of EV biogenesis.

A major difference we observed between EVs secreted by GM1-treated cells and those from control cells was that the former were significantly enriched with GM1, mirroring a similar enrichment in their cells of origin (Fig. 2e).

### Inhibition of ganglioside synthesis impairs the ability of cells to secrete EVs

To investigate the role of endogenously synthesized gangliosides and shed light on the potential implications of decreased ganglioside levels in neurodegenerative diseases, we used three different models of cellular ganglioside reduction: i) N2a cells treated with Genz-123346, a pharmacological inhibitor of glucosylceramide synthase^65^, the first enzyme committed to the synthesis of gangliosides (Fig. 3a). This treatment only partially depletes gangliosides and thus mimics the partial reduction of gangliosides that is observed in HD^36,66^ and other pathophysiological conditions^67–69^; ii) N2a cells overexpressing an N-terminal (exon 1) fragment of mutant huntingtin (mHTT, N2a 97Q cells), a HD model that recapitulates the decrease in GM1 cellular levels described in other HD models and patients’ cells^39,40^; iii) knockout of *B4galnt1* in N2a cells by CRISPR/Cas9^70^, to block the synthesis of complex gangliosides (Fig. 3a) and model a defect of ganglioside biosynthesis in humans^41,42^. As expected, we observed significantly lower levels of GM1 in N2a 97Q cells compared to wild-type N2a cells, given that expression of mHTT decreases ganglioside levels across multiple animal and cell models^36,40^. GM1 levels were further lowered in cells of both genotypes by treatment with 1 µM Genz-123346 (Fig. 3b). Concomitantly, N2a 97Q cells secreted fewer EVs than wild-type N2a cells (Fig. 3c). A further impairment in the secretion of EVs by cells of both genotypes was observed upon inhibition of ganglioside synthesis with Genz-123346 (Fig. 3d and Supplementary Figs. 3a,b), revealing the existence of a positive correlation between EV secretion and intracellular levels of GM1 (Fig. 3e). In line with these conclusions, blocking GM1 synthesis by knockout of *B4galnt1* (Fig. 3f) resulted in a >50% decrease in EV secretion compared to control cells (Fig. 3g and Supplementary Figs. 3c,d).

**Figure 3.**
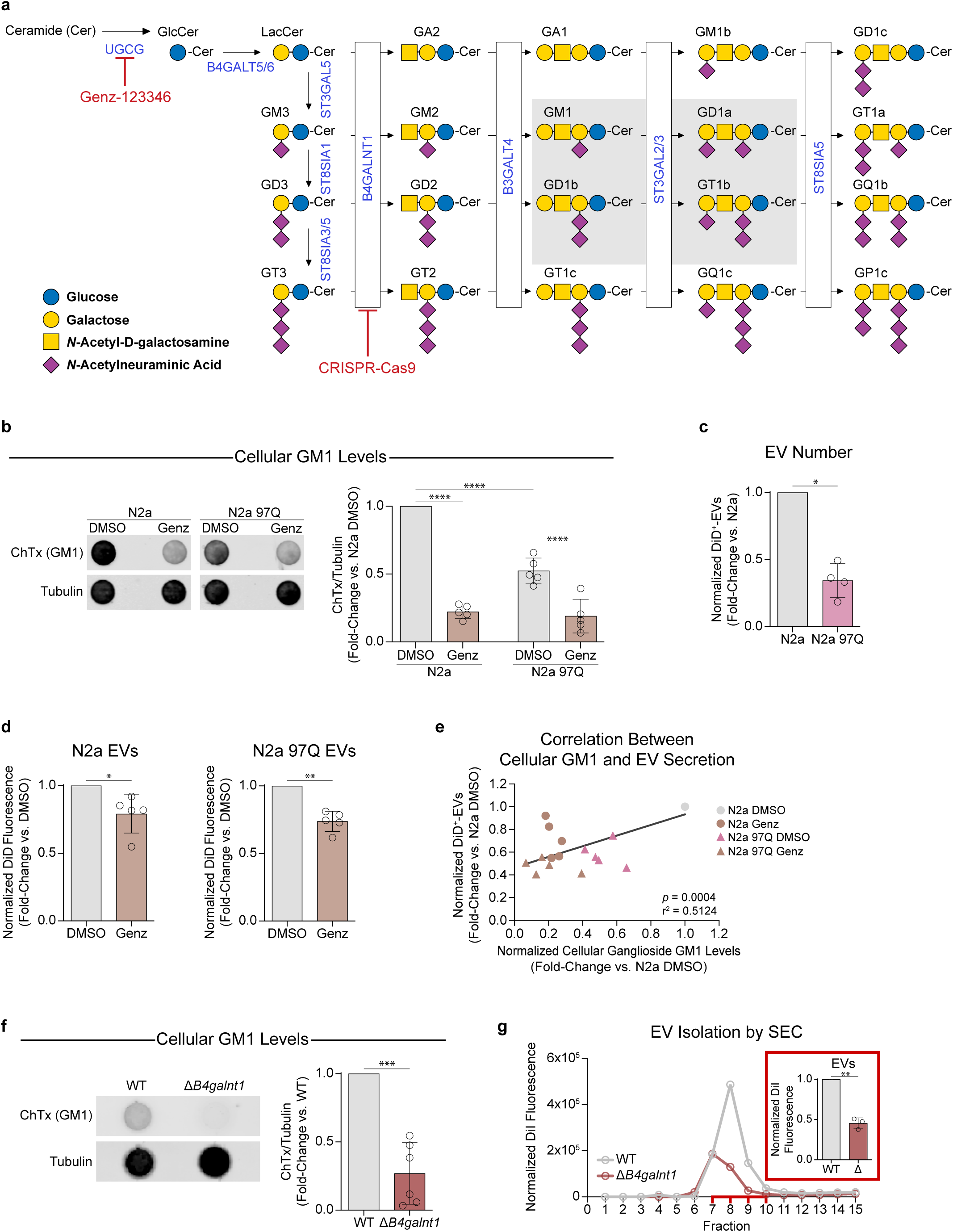
Lowered cellular ganglioside levels result in attenuated EV secretion. **a**. The ganglioside biosynthetic pathway. The glycan moiety of each ganglioside is represented by the Symbol Nomenclature for Glycans (SNFG). Enzymes are indicated in blue text; the four major brain gangliosides are shown in the gray box. Genz-123346 inhibits glucosylceramide synthase, the gene product of *Ugcg*. CRISPR-Cas9 was used to knockout *B4galnt1*. (Modified from Sipione *et al.*, Front. Neurosci., 2020). **b.** Representative dot-blot and densitometric analysis of cholera toxin binding to N2a and N2a 97Q cell lysates to measure GM1 levels after 48 h of treatment with 1 µM Genz-123346. Tubulin was probed as a loading control (n=5 independent experiments). **c.** Fold-change of DiD^+^-EVs secretion by mHTT-expressing N2a 97Q compared to N2a cells, measured by IFC. Data were normalized to cellular protein content (n=4 independent experiments). **d.** DiD^+^-EV fluorescence measured in the CCM of N2a (left) and N2a 97Q (right) cells after cell ganglioside depletion with 1 µM Genz for 48 h. Data were normalized to cellular protein content (n=5 independent experiments). **e.** Pearson correlation between cellular GM1 levels (quantified by a cholera toxin binding assay) and EV secretion (measured by IFC) in N2a and N2a 97Q cells treated with DMSO (vehicle) or Genz-123346 for 48 h. DiD^+^-EV number was normalized to total cellular DiD fluorescence, while GM1 levels were normalized to tubulin (n=5 independent experiments). **f.** Representative dot-blot and relative densitometric analysis of cholera toxin binding to measure levels of GM1 in N2a and N2a Δ*B4galnt1* cell lysates after culture for 22-24 h in serum-free medium supplemented with N-2. Tubulin was probed as a loading control (n=6 independent experiments). **g.** Representative size-exclusion chromatogram of EVs secreted by N2a and N2a Δ*B4galnt1* cells. The DiI fluorescence in each fraction was normalized to cellular DiI fluorescence, and the peaks represent EV-rich fractions (7-10). The inset shows the number of DiI^+^-EVs in the EV-rich fractions measured by IFC and normalized to total cellular DiI fluorescence (n=3 independent experiments). Bars indicate mean ± SD. \**p*<0.05, \*\**p*<0.01, \*\*\**p*<0.001, \*\*\*\**p<*0.0001 by two-way ANOVA with Tukey’s post-hoc test (**b**) or two-tailed paired *t*-test (**c, d, f, g**).

### Administration of exogenous gangliosides restores EV secretion defects in *B4galnt1* KO cells and in HD cells

Reduced EV secretion in *B4galnt1* KO cells could be restored by administration of an equimolar mix (17.5 µM each) of the four major gangliosides (GM1, GD1a, GD1b and GT1b) that are downstream of the synthetic block in *B4galnt1* KO cells, or even 50 µM GM1 alone (Fig. 4a).

**Figure 4.**
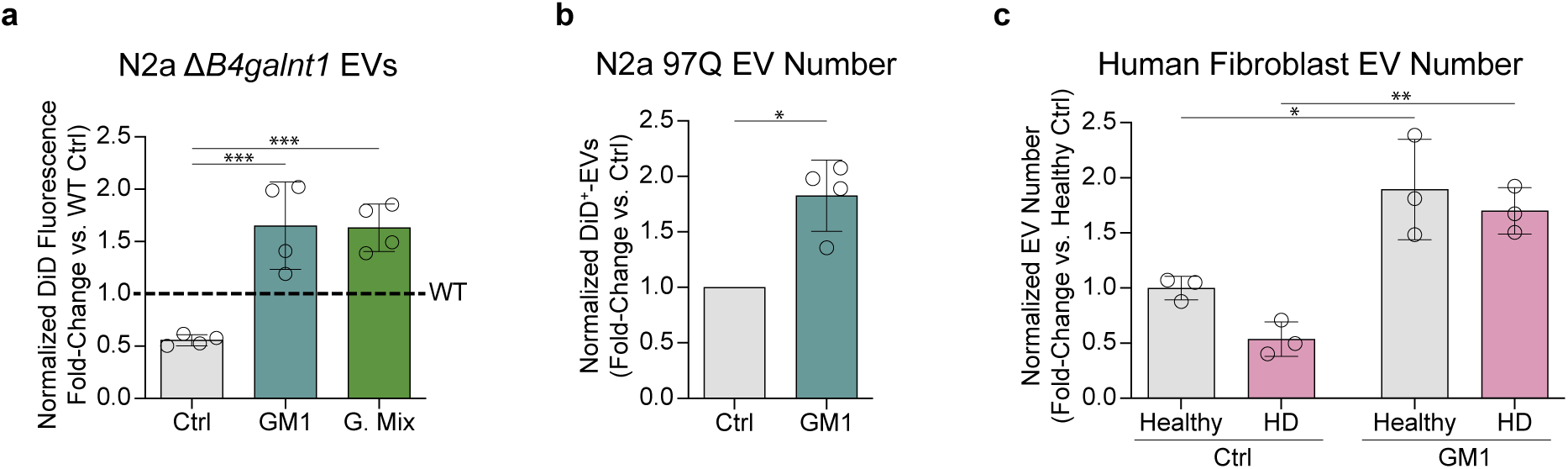
GM1 administration rescues EV secretion in cells with impaired ganglioside synthesis. **a**. Total EV fluorescence in the CCM of N2a Δ*B4galnt1* cells incubated with 50 µM GM1 or a 50 µM equimolar mixture of the major brain gangliosides (G. Mix: GM1, GD1a, GD1b and GT1b) for 22 h (n=4 independent experiments). **b.** Number of DiD^+^-EVs secreted by N2a 97Q cells treated with 50 µM GM1 for 6 h compared to untreated cells (n=4 independent experiments). **c.** Number of EVs secreted by primary human fibroblasts from 3 healthy controls and 3 age-matched HD patients incubated 50 µM GM1 for 22 h, compared to untreated controls. EVs were captured and analyzed using the Exoview platform. Circles represent the data from different donors. EV fluorescence and numbers were normalized to total cellular protein content. Bars indicate mean ± SD. \**p*<0.05, \*\**p*<0.01, \*\*\**p*<0.001 by one-way ANOVA with Tukey’s post-hoc test (**a**), two-tailed paired *t*-test (**b**) or two-way ANOVA with Tukey’s post-hoc test (**c**).

Akin to the rescue of EV secretion in *B4galnt1* KO cells, supplementation of N2a 97Q cells with GM1 resulted in a doubling of EVs secreted in the culture medium (Fig. 4b). To further investigate the defect of EV secretion in HD models and whether it can be rescued by ganglioside administration, we analyzed EV secretion in primary human fibroblasts from three healthy controls and three age-matched HD carriers. All three HD lines -previously shown to have lower GM1 levels than control fibroblasts^40^ – secreted less EVs than healthy fibroblasts (unpaired two-tailed *t*-test, *p*=0.0133), recapitulating the correlation between lowered GM1 levels and impaired EV secretion described for N2a 97Q cells. This reduction was driven by decreased secretion of CD81^+^ and CD63^+^/CD9^+^ double-positive EVs (Supplementary Fig. 4). Administration of GM1 increased EV secretion from all lines and restored EV secretion in HD cells to control fibroblast levels (Fig. 4c). GM1 treatment did not alter the size of EVs secreted by N2a 97Q or HD fibroblasts, in line with our results in WT cells (Supplementary Fig. 2b-d). Altogether, our findings suggest that cell gangliosides are novel key modulators of EV secretion, with decreased levels or lack of gangliosides resulting in impaired EV secretion in disease models, and supra-physiological levels of GM1 (as obtained upon GM1 administration) promoting EV secretion and rescuing EV biogenesis defects.

### Secretion of misfolded proteins within EVs is affected by the cellular ganglioside content

Misfolded proteins, including pathogenic proteins such as mHTT, can be secreted as EV cargo to alleviate cell proteotoxic stress^8,71^. Therefore, defects in EV secretion could affect the ability of cells to dispose of misfolded proteins via EVs. Indeed, N2a 97Q treated with 1 µM Genz-123346 cleared less mHTT via EVs compared to untreated cells, as evidenced by the amount of mHTT in EV fractions isolated by SEC, measured by immunoblotting (Fig. 5a) and ELISA (Fig. 5b). In contrast, cell treatment with GM1 promoted the release of mHTT with EVs (Fig. 5c-d). The mHTT detected in EV fractions was located within the EV particles, similar to ALIX, a classic intralumenal EV marker. This was determined in a proteinase K protection assay that showed that mHTT (and ALIX) could only be degraded by the protease after EV membrane permeabilization with a detergent, to allow the enzyme to reach intravesicular cargo (Supplementary Fig. 5). This suggests mHTT is in the EV lumen, not on the EV surface.

**Figure 5.**
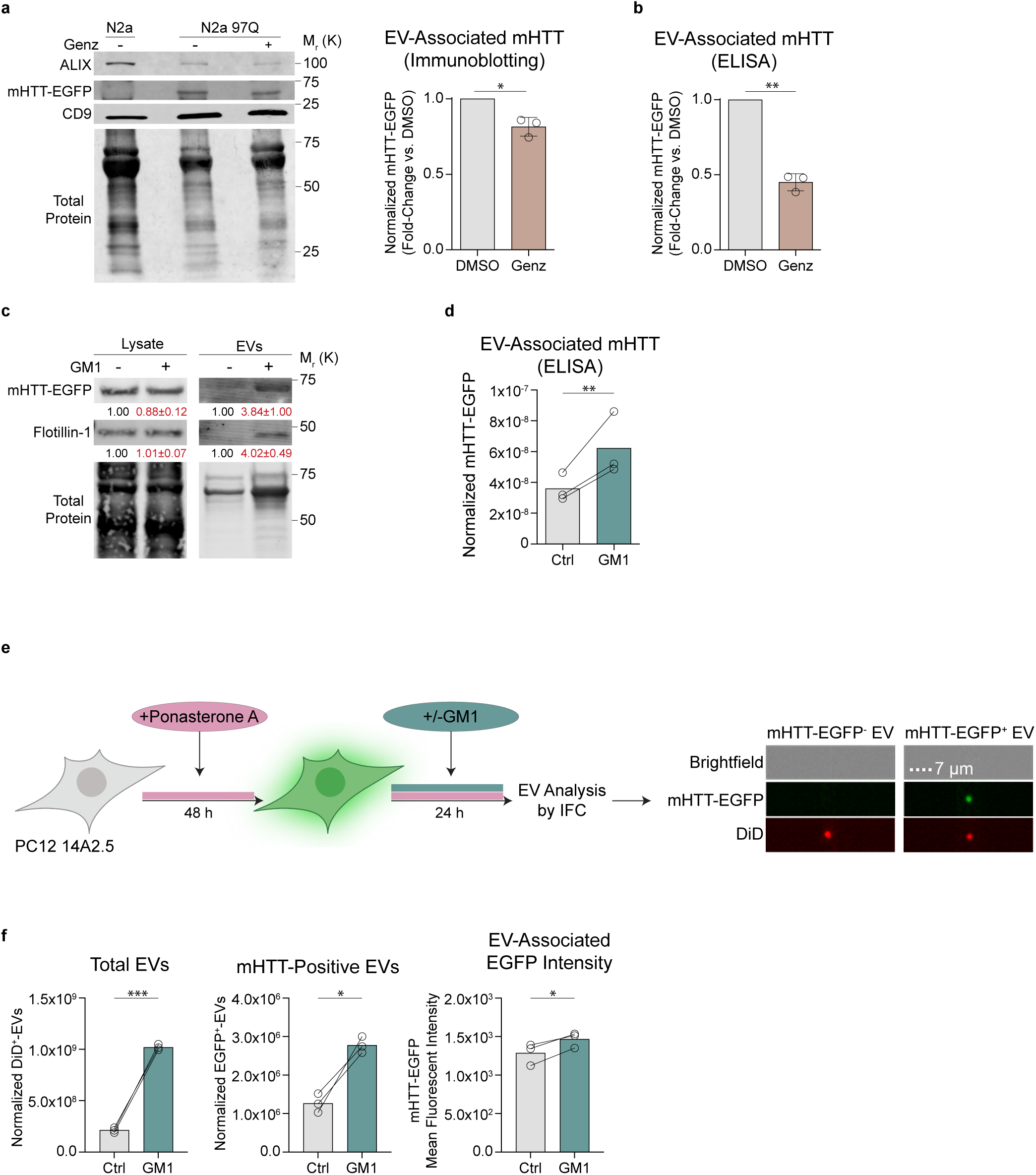
Secretion of mHTT within EVs is modulated by the cellular ganglioside content. **a**. (Left) Representative immunoblots of EVs isolated by size-exclusion chromatography from the conditioned medium of N2a 97Q cells that were treated or not with 1 μM Genz-123346 for 48 h. EVs were collected in the last 24h of incubation with Genz-123346. ALIX and CD9 are EV markers. EV-associated mHTT-EGFP is quantified by densitometric analysis in the graph on the right (n=3 independent experiments). **b.** N2a 97Q cells were treated as in **a.** EV-associated mHTT-EGFP was quantified by ELISA and normalized to total cellular protein content (n=3 independent experiments). **c.** Representative immunoblot for mHTT-EGFP in N2a 97Q cell lysates and EVs isolated from the cleared conditioned medium by ultracentrifugation after cell treatment with 50 µM GM1 for 18h, washing, and EV collection for 24h. Flotillin-1 was probed as an EV marker. The numbers under each band are the mean densitometric values ± SD. Data were normalized over total cellular protein content for EVs (n=2 independent experiments) or over total protein stain for cell lysates (n=3 independent experiments) and are expressed as fold-change vs. untreated controls. **d.** ELISA quantification of mHTT-EGFP in EVs secreted by N2a 97Q incubated for 22h with 50 µM GM1 and isolated by size-exclusion chromatography. Data were normalized to total cellular protein content (n=3 independent experiments). **e.** (Left) Schematic experimental design. PC12 14A2.5 cells were induced with ponasterone A for 48 h to express mHTT-EGFP prior to treatment with 50 μM GM1 for 24 h in the presence of ponasterone A. EVs in the CCM were analyzed by IFC. (Right) Representative IFC images of mHTT-EGFP^-^- and mHTT-EGFP^+^-EVs. **f.** Quantitation of total EVs (DiD^+^, left), EVs carrying mHTT (DiD^+^/EGFP^+^, middle) and the mean EGFP fluorescence intensity of the latter (right) by IFC (n=3 independent experiments). The number of total and mHTT-carrying EVs was normalized to total cellular protein content. Bars and circles show mean ± SD. \**p*<0.05, \*\**p*<0.01, \*\*\**p*<0.001 by two-tailed paired *t*-test (**a, b, f**) or ratio paired *t*-test (**d**).

To determine whether the overall increase in mHTT levels in EV fractions following GM1 treatment was due to enhanced mHTT loading into individual particles or a higher number of EVs carrying the protein, we performed IFC analysis of EVs secreted by PC12 cells that express mHTT-EGFP upon induction with ponasterone A^72^. These cells express higher levels of mHTT-EGFP compared to N2a 97Q, allowing for the specific detection of EGFP^+^-EVs (carrying mHTT-EGFP) and for the relative quantification of EGFP fluorescence within each vesicle by IFC (Fig. 5e). GM1 treatment not only increased the number of EGFP^+^-EVs released in the medium but also increased their mean EGFP fluorescence intensity (Fig. 5f), suggesting that GM1 enhances both the loading of mHTT into individual EVs and the number of EVs carrying mHTT. Furthermore, this increase in the release of mHTT via EVs correlated with a lower intracellular mHTT burden, as shown by the accelerated clearance of intracellular mHTT over time (Fig. 6a-c). Total protein levels were not affected by GM1 (Fig. 5d), suggesting that the observed decrease in mHTT levels was not due to cell death or GM1 effects on general proteostasis. Decreased intracellular mHTT levels were also observed upon GM1 incubation with STHDh^111/111^ cells (Fig. 6e), a striatal HD knock-in model that expresses full-length mHTT from the endogenous *Htt* locus^73^.

**Figure 6.**
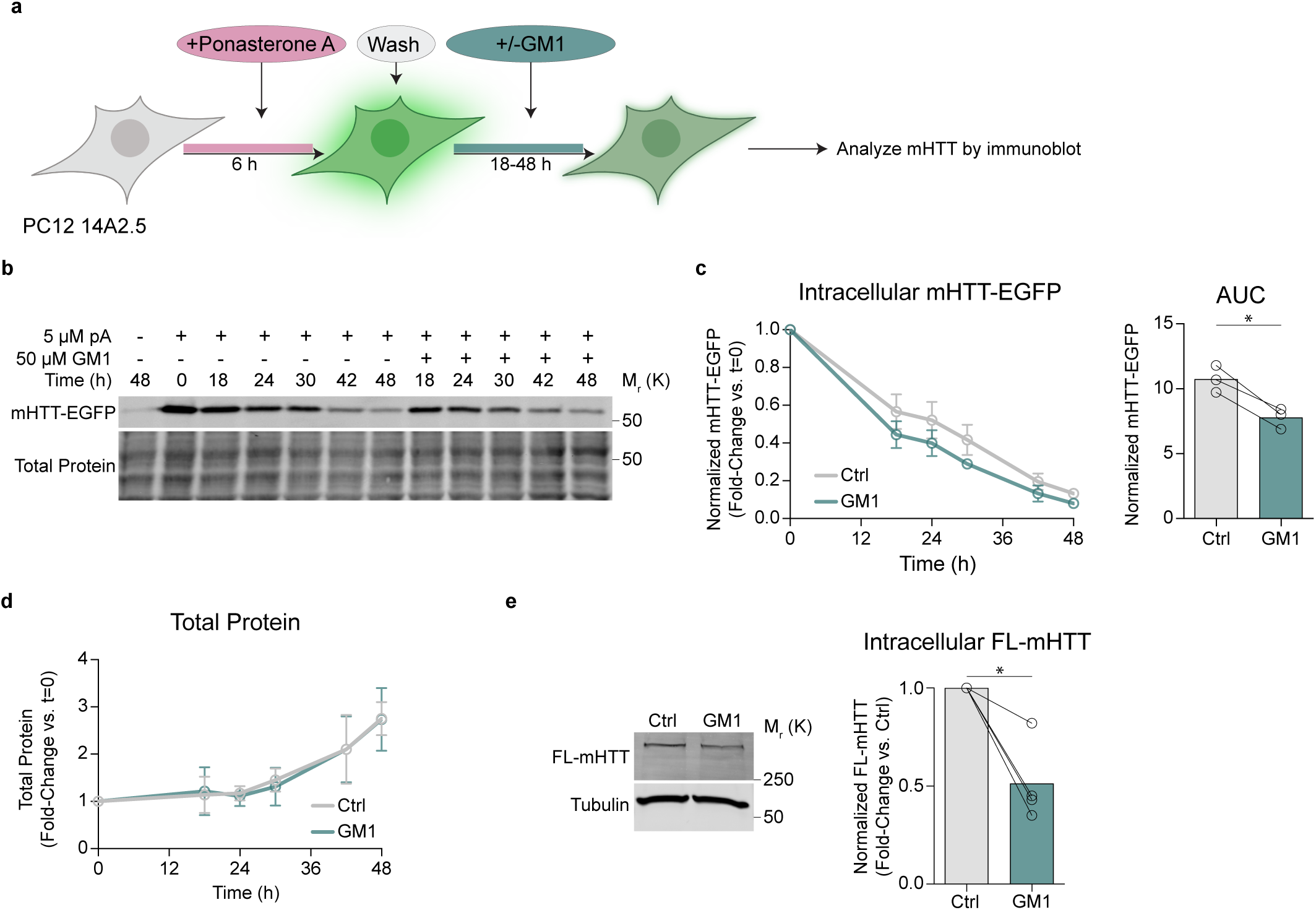
Intracellular mHTT levels are reduced following GM1 treatment. **a**. PC12 14A2.5 cells were induced to express mHTT-EGFP with ponasterone A for 6h. Ponasterone A was then washed-off to allow to follow mHTT clearance over time in untreated cells and cells incubated with 50 μM GM1 for up to 48 h. **b.** Representative immunoblot of the time course of mHTT-EGFP clearance after withdrawal of ponasterone A. **c.** Densitometric analysis of mHTT-EGFP levels normalized to total protein stain and quantitation of cumulative mHTT-EGFP burden (area under the curve, AUC) (n=3 independent experiments). **d.** Total protein quantification in lysates of Ctrl and GM1 treated PC12 14A2.5 cells over 48 h (n=3 independent experiments). **e.** Representative immunoblot and densitometric analysis of intracellular full-length (FL) mHTT in STHDh Q111/111 cells following treatment with 50 μM GM1 for 48 h compared to untreated cells (n=4 independent experiments). Bars and circles show mean ± SD. \**p*<0.05, \*\**p*<0.01, \*\*\**p*<0.001 by two-tailed paired *t*-test.

The ability of GM1 to promote protein clearance via EVs was not limited to mHTT, but extended to other disease models and pathogenic proteins, including N2a cells expressing A53T α-synuclein and HEK cells expressing either wild-type Tau or its disease-associated variants N297K and P301L (Fig. 7). Altogether, our data suggest that through modulation of EV release, gangliosides can affect the disposal of pathogenic proteins via EVs and overall intracellular misfolded protein burden.

**Figure 7.**
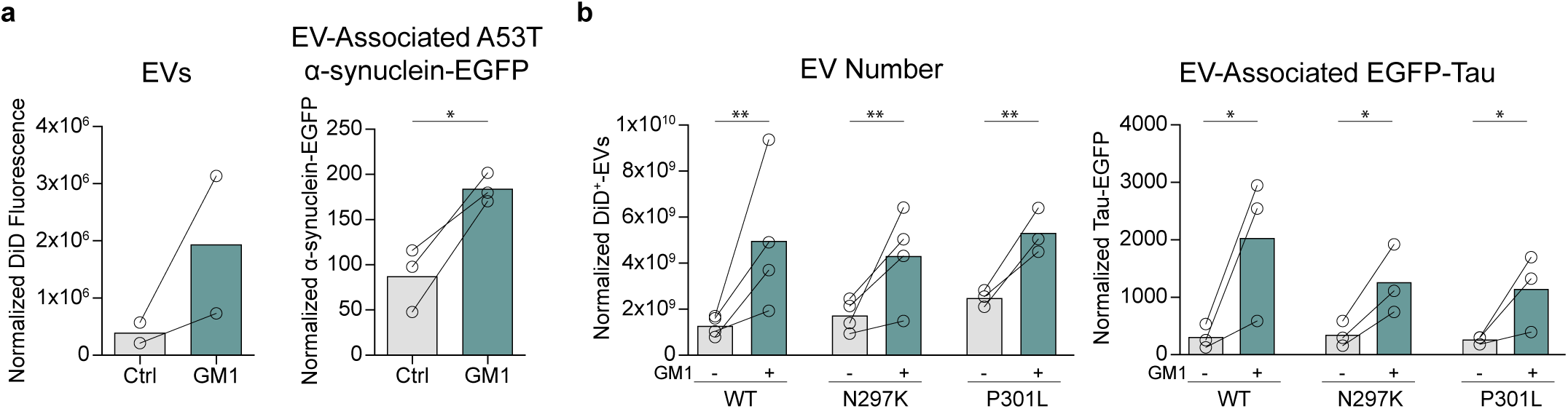
GM1 promotes cellular secretion of alpha-synuclein and tau through EVs. **a**. N2a cells expressing A53T α-synuclein-EGFP were incubated with 50 µM GM1 for 22 h in phenol red-serum-free medium supplemented with N-2. EVs were isolated by size-exclusion chromatography and quantified by fluorometry (left). EV-associated A53T α-synuclein-EGFP was measured by ELISA (right). Data were normalized to total cellular protein content (n=3 independent experiments). **b.** WT GFP-Tau, N279K GFP-Tau or P301L GFP-Tau expression was induced in HEK293T cells with 10ng/mL doxycycline for 72 h. Cells were then incubated with 50 µM GM1 for 22 h EVs were isolated by size-exclusion chromatography and quantified by IFC (left). EV-associated GFP-Tau was measured by ELISA (right). Data were normalized to total cellular protein content (n=3 independent experiments). Bars show means. \**p*<0.05, \*\**p*<0.01 by paired *t*-test (**a**) or two-way ANOVA with Tukey’s post-hoc test (**b**).

### Sialic acid, N-Acetyl-D-Galactosamine and ceramide are key molecular determinants of the modulatory effects of gangliosides on EV secretion

Our studies on EV secretion have so far focused on the effects of a specific ganglioside, GM1, or pharmacological and genetic interventions that broadly reduce the levels of most or all gangliosides, including GM1. While all gangliosides share a common structure - a ceramide lipid tail attached to a variable glycan headgroup (Fig. 8a) - the composition of the glycan headgroup not only defines individual gangliosides, but also dictates their distinct behaviours, functions, and interactions^74^, which can be remarkably different among gangliosides. Therefore, we aimed to investigate whether the ability of GM1 to promote EV secretion is exclusive or a general feature of all gangliosides, regardless of glycan headgroup composition.

**Figure 8.**
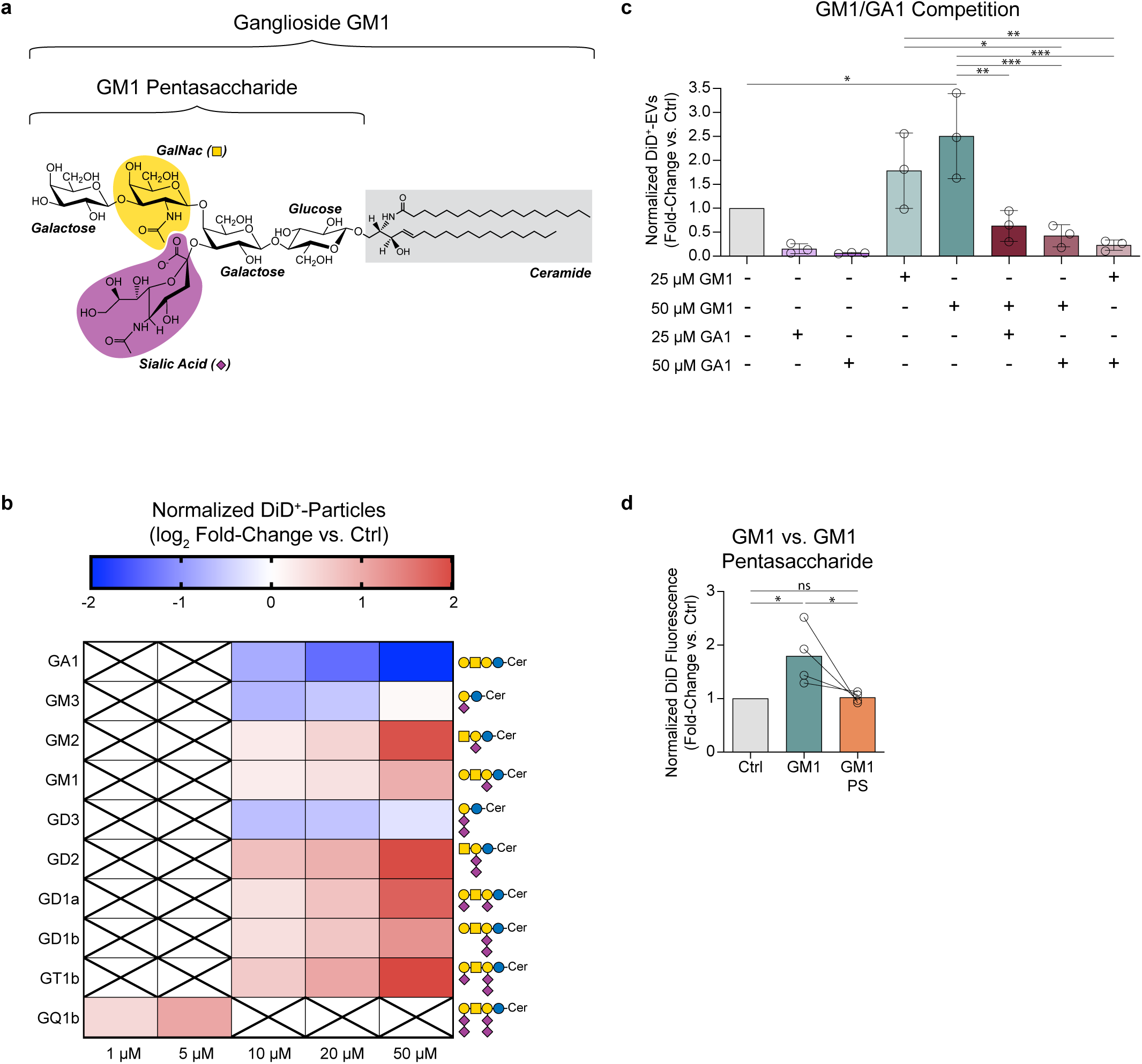
The effects of gangliosides on EV secretion depend on their ceramide tail and the presence of sialic acid and N-acetyl-D-galactosamine in their headgroup. **a**. Structure of ganglioside GM1 as a prototypical ganglioside. N-Acetyneuraminic acid (sialic acid), N-acetyl-D-galactosamine (GalNac) and the ceramide tail are highlighted in purple, yellow and gray, respectively. The SNFG symbols for sialic acid and GalNac are shown in brackets beside the sugar residue name. b. Heatmap of the log_2_ fold-change of the number of EVs secreted from N2a cells treated with different gangliosides at the indicated concentrations for 22 h compared to their respective controls. Blue shades indicate decreased EV secretion, red shades indicate increased EV secretion. The SNFG structures of each tested ganglioside is shown on the right of the heatmap. The number of EVs in the CCM was measured by IFC and normalized to total cellular protein content (n=3-5 independent experiments). All treatments significantly affected EV secretion (*p*<0.05 by Friedman’s test). c. N2a cells were treated with ganglioside GM1 or GA1 individually or together at the indicated concentrations for 22 h in phenol red-serum-free medium supplemented with N-2. The number of EVs in the CCM was measured by IFC and normalized to total cellular protein content (n=3 independent experiments). d. Fluorescence of EVs secreted by N2a cells treated with 50 µM GM1 or GM1 pentasaccharide (GM1 PS) for 22 h. EV fluorescence was normalized to total cellular protein (n=5 independent experiments). Heatmap cells are means. Bars represent means ± SD. \**p*<0.05, \*\**p*<0.01, \*\*\**p*<0.001 by Friedman’s test (**b**), repeated measures one-way ANOVA with Tukey’s post-hoc test (**c**) or one-way ANOVA with Tukey’s post-hoc (**d**).

To address this question, we tested the effects of all major gangliosides on EV secretion. All complex gangliosides stimulated EV secretion in N2a cells, with those carrying more sialic acid residues, including GD2, GD1a, GD1b and GT1b, being more effective, and GQ1b being more potent than GM1 (Fig. 8b). In contrast, the simpler gangliosides GM3 and GD3 inhibited EV secretion. Strikingly, ganglioside GA1 (asialo-GM1), which shares the glycan structure of GM1 but lacks the sialic acid residue, was a very effective inhibitor of EV secretion (Fig 8b). Furthermore, GA1 abrogated the EV-promoting effects of GM1, reducing EV secretion to very low levels in both GM1-treated and untreated N2a cells (Fig 8c). These findings demonstrate that the ability to promote EV secretion is not a universal property of all gangliosides and depends on specific ganglioside glycan structures. In particular, the distinct behaviours of simple (GM3 and GD3, which inhibit EV secretion) versus complex gangliosides (which promote EV secretion) suggest that the presence of the N-acetyl-D-galactosamine (GalNac) residue in complex gangliosides may be a critical determinant of the EV-promoting effects of gangliosides. The presence of at least one sialic acid residue also appears essential, as its absence reverses the ganglioside effects on EV secretion.

Gangliosides localize to membrane microdomains, and their functions often require or are mediated by their ceramide tail^33^. To determine whether this is also true for their EV-promoting activity, we treated N2a cells with either GM1 or the GM1 pentasaccharide, which shares the same glycan structure of GM1 but lacks its ceramide moiety. The pentasaccharide was unable to recapitulate the effects of GM1 on EV secretion (Fig. 8d), suggesting that membrane anchoring via the ceramide tail is necessary for gangliosides to modulate EV secretion.

## Discussion

Gangliosides are multifunctional molecules with numerous roles in both health and disease^29,30^. Here, we have identified a novel function of GM1 and other gangliosides in modulating EV secretion. The modulatory effects of gangliosides on EV biogenesis may have significant implications in neurodegenerative diseases where levels of gangliosides are altered, and may help explain some of the previously described therapeutic effects of GM1 in models of misfolded protein diseases^45,47,49^.

Using several complementary approaches, we have shown that GM1 administration to neuroblastoma N2a cells stimulates EV secretion. This effect is further enhanced with continuous administration of the ganglioside. To minimize sample loss and avoid biases introduced by EV isolation methods that rely on specific markers – the expression of which may vary across cell types and treatments - we initially used direct IFC analysis of in-cell fluorescently labelled EVs^53^. We then confirmed our findings by quantifying EVs isolated by ultracentrifugation or size-exclusion chromatography, the two most widely used EV isolation methods^55,75^. The GM1-induced increase in the number of EVs secreted was mirrored by a rise in total protein and EV markers in EV preparations from both N2a cells and primary rat neurons, confirming that we measured *bona fide* EVs.

GM1 significantly increases EV secretion not only in immortalized and primary neuronal cells from mice and rats, but also in human cell lines (HeLa and HEK293 cells), primary human skin fibroblasts and human iPSC-derived neurons. The ability of GM1 to increase EV secretion across species, cell types, or tissue origins suggests a fundamental role for gangliosides in EV production. In contrast, disruption of endogenous ganglioside biosynthesis through pharmacological (Genz-123346) or genetic (CRISPR-Cas9 KO of *B4galnt1*) approaches significantly decreases EV secretion, with cellular GM1 levels positively correlating with the amount of secreted EVs. This finding suggests that it is not only the supra-physiological cellular ganglioside levels achieved by pharmacological administration of GM1 that promotes EV secretion, but rather endogenous gangliosides may play an integral role in constitutive EV secretion. Supporting these conclusions, knock out of *B4galnt1* (encoding GM2/GD2 synthase) in N2a cells significantly decreased EV secretion, whereas in another study, overexpression of *B3GALT4* (encoding GM1 synthase) in a breast cancer cell line increased levels of the EV markers TSG101 and CD63 in EV fractions, suggesting enhanced EV release^76^.

Whether GM1 treatment preferentially enhances the production of exosomes or ectosomes remains to be determined. We found that GM1 did not specifically increase secretion of small or large EVs as EV size distribution remained unchanged across the various cell types analyzed. Some studies have attempted to differentiate EVs (exosome vs. ectosome) based on their tetraspanin profiles. While CD63 has been proposed as a specific marker for exosomes^56^, recent findings in HeLa cells suggest a more complex relationship between EV origin and tetraspanin profile, with CD63, CD9 and CD81 all transiently localizing to both multivesicular bodies and the plasma membrane following translation, ruling out the presence of any single tetraspanin on a vesicle as a marker of its origin^77^. Regardless of the baseline tetraspanin profile of EVs secreted by different cell types – which was different from one cell type to another - GM1 treatment proportionately increased all EV subtypes, suggesting a general mechanism of action rather than selective modulation of a specific EV subpopulation.

To investigate the structural requirements underlying the EV-promoting activity of GM1, we examined how different ganglioside features influence EV secretion. Both the specific composition of the glycan headgroup and the presence of the ceramide lipid tail proved essential. The GM1 pentasaccharide alone failed to enhance EV secretion, implying that ganglioside incorporation into membranes through their ceramide tail is required. The integration of gangliosides into cell membranes dictates their intracellular traffic, subcellular localization, and affects membrane properties and curvature^33,78^. Of note, GM1 and other gangliosides induce positive membrane curvature^33,34^, which might promote EV biogenesis both at the plasma membrane and in MVB through the formation of intraluminal vesicles, similar to the well-established effects of ceramide on exosome biogenesis^79^. However, membrane curvature alone is unlikely to fully explain the effects of gangliosides on EV secretion. Catabolism of gangliosides to EV-promoting lipids like ceramide and sphingosine^80,81^ is also an unlikely mechanism. While most gangliosides induce positive membrane curvature, although to different extents^33^, and all gangliosides are catabolized to ceramide and sphingosine, our data clearly show that the specific glycan structures modulate both the magnitude and direction of their effect on EV secretion. This strongly suggests that gangliosides influence EV secretion through the molecular interactions of their glycan headgroups, rather than via their shared sphingolipid catabolites.

Notably, gangliosides with higher sialic acid content – typically associated with greater membrane curvature and stronger segregation properties in lipid bilayers^33^ - were more effective than GM1 in promoting EV secretion. In contrast, GM3 and GD3 inhibited EV release. Additionally, GA1, a ganglioside structurally similar to GM1 but lacking sialic acid, strongly inhibited EV secretion in a dose-dependent manner. Furthermore, co-administration of GA1 and GM1 abolished the EV-promoting effect of GM1, resembling a dominant-negative interaction. Since GA1 induces positive membrane curvature to a degree comparable to GM1^82^ yet suppresses EV secretion, our findings further support the conclusion that the effects of gangliosides on EV biogenesis are mainly driven by specific sugar residues rather than membrane curvature alone. The inhibitory effects of GM3 and GD3, which lack GalNAc in their headgroup, suggest that both sialic acid and GalNAc residues are required for the positive effects of gangliosides on EV secretion. We propose that gangliosides may engage protein partners involved in EV biogenesis via these residues, and that partial engagement with only one of these sugars might hamper EV biogenesis. Alternatively, intramolecular interactions between the sialic acid and the GalNAc residue, which confer a more rigid conformation to the headgroup of complex gangliosides^33^, may be necessary for the interaction with partners in EV biogenesis and their functional effects.

Many proteins directly implicated in EV biogenesis, including ESCRT components, flotillin-1, syndecan-1, and tetraspanins, localize to membrane microdomains enriched with gangliosides^23–27,83^. A recent study using a clickable photoaffinity GM1 probe identified CD9 and other proteins involved in EV biogenesis as part of the human GM1 interactome^84^. Therefore, interactions between gangliosides and EV biogenesis proteins may underlie their modulatory effects on EV secretion, by stabilizing or sequestering factors in lipid rafts, altering their conformation and activity, or modulating their interactions with partner proteins^85–89^. Thus, gangliosides may fine-tune vesicle formation and release by shaping the molecular environment where EV biogenesis occurs.

Our finding raises important questions about how EVs are affected by ageing, when brain levels of complex gangliosides decline^90^, and in pathologies where ganglioside metabolism is disrupted. The N2a *B4galnt1* KO cells used in our study model the ganglioside biosynthetic defect resulting from mutations in the human B4GALNT1 gene, which causes a form of hereditary spastic paraplegia (HSP)^41^. The impaired EV secretion observed in these cells may reflect a similar defect in patient-derived cells and contribute to disease pathogenesis, given the role of EVs in processes that are key to normal brain function, including development, synaptic plasticity, proteostasis and microglia activity^5–7^.

In HD, mHTT expression reduces cellular and brain levels of GM1 and other major gangliosides^36,40^, a finding replicated in the HD cell models used in this study. We show that HD cells (N2a 97Q and primary fibroblasts) secrete fewer EVs than controls, a phenotype reversed by GM1 administration. Our data represent one of the first experimental confirmations^91^ of impaired EV secretion in HD, and support previous computational predictions^14^. However, they are in contrast with another study reporting increased EV secretion from HD fibroblasts and derived neural stem cells^92^. The discrepancy may stem from differences in EV isolation and analytical methods used, as Beatriz *et al.* focused their analysis on small EVs isolated by ultracentrifugation^92^, a method that may result in EV losses^53,93^ and might bias results, while our approach minimizes sample loss and captures a broader EV population. Although multiple mechanisms might be responsible for the changes in EV secretion observed in HD cells^91^, our data suggest that the reduction in ganglioside levels is key, since the phenotype is rescued by GM1 administration.

EVs carrying misfolded proteins play important roles in brain proteostasis^8,71,94^, reducing the intracellular accumulation of pathogenic proteins and mitigating their toxicity^8,9,71^. Thus, impaired EV production in HD cells could have a negative impact not only on brain processes modulated by EVs, but also on the cells’ ability to export and clear mHTT through this pathway. Our studies support this hypothesis, by showing that pharmacological inhibition of ganglioside synthesis with Genz-123346 decreases the amount of mHTT secreted via EVs. In contrast, GM1 treatment increases the secretion of EV-associated mHTT in a neuronal model and human HD fibroblasts. A proteinase K protection assay allowed us to determine that mHTT is loaded into EVs as a luminal cargo, rather than associated to their surface. In an inducible PC12 cell model, GM1 treatment increases both the number of mHTT-containing EVs released in the medium and the amount of mHTT loaded into each particle, while concomitantly decreasing the intracellular mHTT burden. These results highlight the therapeutic potential of gangliosides and align with and contribute to explain the mHTT-lowering effects of GM1 *in vivo* in HD mouse model, which result in neuroprotection and restoration of normal motor and non-motor behaviour^45,46^. Given the greater effectiveness of other gangliosides (GD2, GD1a, GD1b and GT1b) compared to GM1 in promoting EV secretion, the question arises whether they might confer even stronger neuroprotective effects. However, GM1’s overall neuroprotective effects might include a range of mechanisms beyond EV secretion^95^, which might not be shared by other gangliosides.

Beyond HD, GM1 promotes EV-mediated secretion of A53T α-synuclein and wild-type and mutant tau - pathogenic proteins implicated in familial PD, AD and tauopathies, respectively – suggesting broader therapeutic potential. Further studies are needed to elucidate the mechanism by which gangliosides influence misfolded protein packaging into EVs and determine whether specific chaperones facilitate this process. The cysteine-string protein (CSPα, or DnaJC5) was shown to play a role in mHTT loading into 180-240 nm and larger vesicles^96^, but whether it is involved in the effects of GM1 described in our studies remains to be determined.

A potential concern regarding the use of gangliosides to enhance misfolded protein secretion via EVs is whether this could promote the prion-like spreading of pathogenic proteins throughout the brain^10,12^. However, this appears unlikely, given strong experimental evidence from *in vivo* models demonstrating that GM1 administration slows or even halts neurodegeneration, lowering brain levels of mHTT and α-synuclein^45,46,49^. We speculate that GM1 not only increases EV production but also modifies their fate. In addition to increasing cellular ganglioside levels, GM1 treatment increases its own levels in secreted EVs. This may have significant consequences for EV fate and function, as gangliosides reside in the outer leaflet of EV membranes and may contribute to recognition, interaction and uptake by target cells. Gangliosides on EVs have been implicated in EV internalization through interactions with integrins and sialic acid-binding siglec receptors on recipient cells in various tissues and cell types^70,97^. In the brain, EV enrichment with GM1 could facilitate uptake and enhance degradation of misfolded protein cargo by microglia, preventing prion-like propagation and explaining why GM1 administration in HD mice reduces brain mHTT levels and mitigates neurodegeneration. Beyond acting as a ligand for EV uptake, EV gangliosides may also exert signaling functions, as seen with EV-associated GD3, which mediates T-cell arrest in the ovarian tumor microenvironment^91^. Given our finding that depletion of cellular gangliosides - either pharmacologically (Genz-123346) or genetically (CRISPR-Cas9 KO of *B4galnt1*) - reduced ganglioside content in secreted EVs, modulating cellular ganglioside levels could represent a method to alter EV-mediated signaling and cargo delivery. Changes in ganglioside expression during development, ageing, or pathological conditions could similarly influence EV biology by affecting their interactions with recipient cells.

A notable finding in our study with implications for cancer therapy is the ability of GA1 to block EV production in neuroblastoma cells. Tumor-derived EVs play critical roles in tumor progression, metastasis and immune evasion^98,99^. Given these key roles and that many tumors upregulate EV biogenesis^100,101^, inhibiting EV secretion with GA1 could provide a novel therapeutic approach, potentially disrupting multiple tumor-supporting processes.

Overall, our study demonstrates that gangliosides, both endogenous and exogenously applied, are potent modulators of EV secretion across multiple cell types. By elucidating their role in EV biogenesis, we provide insight into novel pathogenic mechanisms in diseases characterized by ganglioside deficiency and into how GM1 may exert its mHTT-lowering and neuroprotective effects. Future work should address the precise molecular interactions by which gangliosides influence EV biogenesis and cargo loading, to harness these effects to improve proteostasis and other EV-dependent processes.

## Methods

### Cell models

***Mouse neuroblastoma Neuro2a cells (N2a)*** were kindly donated by Dr. Satyabrata Kar (University of Alberta, Canada). N2a cells were cultured in DMEM:Opti-MEM I (1:1) supplemented with 10% heat inactivated FBS, 2mM L-glutamine and 0.11g/L sodium pyruvate (N2a growth medium) and maintained at 37°C with 5% CO2.

#### Generation of N2a cells stably expressing mHTT-EGFP or α-synuclein (A53T)

N2a cells were stably transfected with the cDNA for mHTT Exon 1 (containing 97 glutamines) tagged with enhanced green fluorescent protein (HTT-Exon1 (97Q) EGFP) kindly donated by Dr. Janice Braun (University of Calgary, Canada), or α-synuclein (A53T) tagged with enhanced green fluorescent protein purchased from Addgene (plasmid #40823). Cell transfection was performed using Lipofectamine 3000® reagent (Thermo Fisher Scientific, USA). Geneticin® (800μg/mL) was added to the medium to select stably transfected cells. GFP-fluorescent cells were further sorted into 96 well-plates using FACS Aria™ III cell sorter (BD Biosciences, USA), at the Faculty of Medicine and Dentistry Flow Cytometry Facility, to select stably transfected clones. Transgene expression in selected clones was confirmed by immunoblotting. Stably expressing cell clones, referred to as N2a 97Q or N2a α-synuclein (A53T), were maintained in N2a growth media with 400μg/mL of Geneticin® at 37°C with 5% CO_2_. Geneticin was removed from the culture media one passage prior to performing experiments. Cells were routinely checked for GFP fluorescence by microscopy and were periodically re-sorted by fluorescence-associated cell sorting (FACS) when the percentage of fluorescent cells fell below 80%.

***N2a* Δ*B4galnt1*** were obtained by deletion of *B4galnt1* from WT N2a cells using CRISPR-Cas9 as previously described^70^, and maintained in N2a growth medium at 37°C with 5% CO2.

***HeLa cells*** were kindly donated by Dr. Ing Swie Goping (University of Alberta, Canada) and grown in DMEM supplemented with 10% heat inactivated FBS, 2mM L-Glutamine and 0.11g/L sodium pyruvate (HeLa growth medium), at 37°C with 5% CO2.

***Embryonic rat cortical neurons*** were isolated from E18 embryos and maintained in neuronal growth media (Neurobasal™ containing 200μM Glutamax®, 100U/mL penicillin, 100μg/mL streptomycin and 1X B27^®^ supplement) on plates coated with poly-L-lysine (molecular weight ≥ 300,000, Sigma Aldrich, USA). All procedures on animals were approved by the University of Alberta’s Animal Use Committee and were in line with the Canadian Council on Animal Care (CCAC). Briefly, pregnant dams (Sprague Dawley) were subjected to isoflurane-anesthesia followed by decapitation. Thereafter, the uterus was transferred to a petri dish on ice and embryos were isolated, decapitated and the brains were dissected under a microscope. Dissected cortical tissue was resuspended in 0.25% trypsin-EDTA (Thermo Fisher Scientific, USA) in a 50-mL polystyrene tube and incubated for 20 min at 37°C in a water bath. The digested tissue was centrifuged at 300 x g for 2 min, the supernatant discarded, and the pellet resuspended in trituration media (DMEM + 10% heat inactivated FBS + 1% penicillin/streptomycin). Thereafter, the tissue was passed 10 times through a fire-polished glass Pasteur pipette and the supernatant obtained after tissue sedimentation was transferred to a new polystyrene tube. Trituration was performed three times in total. The cell suspension was then filtered through a 40 μm pore-size nylon cell strainer (Thermo Fisher Scientific, USA). Cells were counted using a haemocytometer and plated at a density of 0.13x10^6^ cells/cm^2^ in trituration medium. After cells had attached (1.5 to 2 h after seeding), the trituration medium was replaced with neuronal growth media. On DIV2, cells were treated with cytosine arabinoside (3μM, Sigma Aldrich, USA) for 48 h to block the proliferation of glial cells and reduce their presence to less than 1%^102^. Half of the medium was replaced with fresh medium every 4 days. Cultures were maintained at 37°C with 5% CO2. Experiments were performed on DIV 15.

***Primary human fibroblasts***, purchased from the Coriell Institute for Medical Research, USA, were derived from 1 female and 2 male HD donors (HD11M, HD29F, HD35M; Coriell codes GM04855, GM03621, GM04208) and three age-matched healthy male donors (WT15M, WT33M, WT36M; Coriell codes GM01869, AG08181, AG07573). A complete characterization of these cells is available in the Coriell Institute for Medical Research website. Fibroblasts were maintained in MEM (with non-essential amino acids) supplemented with 15% heat inactivated FBS, 2mM L-Glutamine and 0.11g/L sodium pyruvate (fibroblast growth media), at 37°C with 5% CO2. For cell passaging and seeding, cells were washed with HBSS (Ca^2+^/Mg^2+^-free) and then incubated with the same buffer containing 0.53mM EDTA for 2 min at r.t., prior to cell trypsinization in 0.05% trypsin-EDTA (Corning Life Sciences, USA). Cells were passed before they could reach 80% confluence as cellular ganglioside levels may change upon reaching confluence^103^.

#### Differentiation of human iPSCs into i^3^Neurons

Human iPSCs (on a male WTC11 background) expressing mNGN2 under the control of a doxycycline inducible promoter^104^ were cultured on plates coated with Matrigel in KnockOut™ DMEM, grown in Essential 8 medium and maintained at 37°C with 5% CO2. The medium was changed every day and cells were split once they reached 80% confluency using Accutase. To generate glutamatergic neurons (i^3^Neurons), iPSCs were differentiated following a previously described protocol^104^. Briefly, i^3^N iPSCs were pre-differentiated in growth medium composed of KO-DMEM medium supplemented with NEAA, mouse laminin (1 ug/mL), Y-27632-ROCK inhibitor (3.2 mg/mL), BDNF (10 ng/mL), NT3 (10 ng/mL), N-2 supplement and doxycycline (2 µg/ml) for 3 days, with daily medium changes and ROCK inhibitor removed from day 2. Pre-differentiated cells were then dissociated using StemPro Accutase, and seeded in plates coated with high molecular weight poly-D-lysine in differentiation medium composed of DMEM/F12-HEPES:Neurobasal-A medium (1:1), supplemented with NEAA, mouse laminin (1 ug/mL), BDNF (10 ng/mL), NT3 (10 ng/mL), GlutaMax and 0.5× N-2 and 0.5× B27 supplement. Half of the medium was replaced on day 7. On days 14 and 21, half the medium was replaced with double the volume, then one-third of the medium was replaced weekly until i3Neurons were used on day 56.

***PC12 14A2.5*** cells expressing mHTT(exon 1)-EGFP were previously described^72^. They were maintained in DMEM supplemented with 5% heat inactivated FBS, 10% heat inactivated horse serum, 2mM L-Glutamine and 0.11g/L sodium pyruvate, at 37°C with 5% CO2. Cell passaging was performed by trypsinization in 0.05% trypsin-EDTA (Corning Life Sciences, USA). Expression of mHTT-EGFP was induced by incubating cells with 5 µM ponasterone A for 6 or 72 h as indicated, and as previously described^72^.

***STHDh 111/111,*** conditionally immortalized knock-in striatal cells expressing full-length mHTT were maintained in DMEM supplemented with 10% heat inactivated FBS, 2mM L-Glutamine and 0.11g/L sodium pyruvate at 33°C with 5% CO_2_, as previously described^73^ .

***Inducible HEK 293 cells expressing WT, N297K, or P301L tau*** were engineered using the Flp-In T-REx™ HEK293 Cell Line (Thermo Fisher Scientific, USA). These cell lines are characterized by targeted integration of a construct for GFP-0N4R tau (wild-type or mutant) under the control of a doxycycline-inducible promoter into a single FRT integration site within the cell genome. The generation of GFP-0N4R P301L tau cell line has been previously described^105^. The N279K and WT GFP-0N4R tau cell lines were generated by the same procedure. Tau HEK293 cells were cultured in DMEM high glucose supplemented with 10% FBS, glutamine (2 mM), sodium pyruvate (1 mM) and Hygromycin B (0.15 mg/ml), and maintained at 37°C with 5% CO_2_. Cells were treated with 10 ng/ml doxycycline for 72 h to induce WT or mutant GFP-0N4R tau expression.

### In-cell labelling for EV detection

Indirect fluorescent labelling of EVs was performed as previously described^53^ to detect and quantitate EVs by fluorometry or imaging flow cytometry. Cell membranes (including EV membranes) were labelled with the lipophilic membrane stain DiI (1,1’-dioctadecyl-3,3,3’,3’-tetramethylindocarbocyanine perchlorate; λ_Ex_/λ_Em_ = 549/565 nm) or DiD (1,1’-dioctadecyl-3,3,3’,3’-tetramethylindodicarbocyanine, 4-chlorobenzenesulfonate salt; λ_Ex_/λ_Em_ = 644/665 nm), according to the manufacturer’s instructions (Invitrogen™, Thermo Fisher Scientific, USA). Briefly, 5μL of the dye solution (1mM) was added to each 1mL of cell suspension containing 1x10^6^ cells in serum-free media and incubated for 8-20 min in the dark at 37°C in a polypropylene tube. The suspension was then centrifuged at 400 x g for 5 min at r.t. and the stained cell pellet was washed three times in serum-containing growth medium to remove any unbound dye. The cells were then resuspended in growth medium and seeded in 100mm dishes. The seeding densities were determined such that the confluence of the cells at the time of harvesting was approximately 80%.

### Preparation of EV collection medium

The medium used for EV collection from cells was either serum-free medium supplemented with heat inactivated EV-depleted FBS (EVD complete medium) or phenol red-free, serum-free medium supplemented with N-2 supplement (SFM+N-2). To prepare EV-depleted FBS, heat inactivated FBS was transferred to 26.3mL thickwall polycarbonate tubes (Beckman Coulter, USA) and centrifuged at 100,000 x g for 16 h at 4°C, using a Type 70 Ti Fixed-angle rotor (Beckman Coulter, USA) in an Optima L-70 ultracentrifuge (Beckman Coulter, USA)^106^. The supernatant was filter-sterilized with a 0.22μm PVDF filter (Millipore, USA) and stored at -20°C until further use. Collection medium containing N-2 supplement was filtered through a 0.1 µm PVDF filter (Millipore, USA) prior to use.

### Cell treatment with gangliosides

Porcine ganglioside GM1 was dissolved in cell-culture grade DPBS or DMSO to a stock concentration of 10 mM. Bovine gangliosides GM2, GD3, GD2, GD1a, GD1b, GT1b, GQ1b, and the GM1 pentasaccharide were dissolved in cell-culture grade DPBS to a stock concentration of 10 mM. GA1 and GM3 were dissolved in cell-culture grade DMSO to a stock concentration of 10 mM. Gangliosides were added to cells in growth media or SFM+N-2 upon cell seeding or after cell attachment to the growth plate (2-4 h after seeding) at 1-50 µM concentration, as indicated. Where indicated, gangliosides were washed off with two washes with HBSS (with Ca^2+^/Mg^2+^) supplemented with 0.2% essentially fatty acid-free BSA, followed by two washes with HBSS (with Ca^2+^/Mg^2+^).

### Inhibition of ganglioside synthesis with Genz-123346

Genz-123346 was solubilized in 100% cell-culture grade DMSO to a stock concentration of 10 mM and stored at -20°C until use. It was added to cells in growth media at the time of cell seeding to a final concentration of 1μM (the final concentration of DMSO was 0.01%). DMSO at the final concentration of 0.01% was added to vehicle control groups. After 24 h of treatment, cells were washed with HBSS (with Ca^2+^/Mg^2+^) and the treatment was replenished in EV collection media.

### EV collection and isolation

After cell treatment, the medium was replaced with EV collection medium and allowed to be conditioned by cells for 16-24 h to collect EVs. To remove loose cells, apoptotic bodies and cell debris, the conditioned medium was collected in polypropylene tubes and centrifuged at 2,000 x g for 10 min at 4°C in an Eppendorf® Centrifuge 5810 R, using an A-4-62 swinging bucket rotor. The supernatant retained for analysis is hereafter referred to as the cleared conditioned medium (CCM). To avoid EV loss, where possible, pre-lubricated low protein-binding microcentrifuge tubes were used in the subsequent steps of EV isolation and analysis. Depending on the experiment, ultracentrifugation or ultrafiltration followed by size-exclusion chromatography were used to isolate EVs.

#### Ultracentrifugation (UC)

The CCM was ultracentrifuged at 100,000 x g for 90 min at 4°C in a 10.4mL thickwall polycarbonate tube (Beckman Coulter, USA), using an MLA-55 fixed angle rotor (Beckman Coulter, USA) in an Optima™ MAX-XP tabletop ultracentrifuge (Beckman Coulter, USA). The EV pellet was washed by resuspension in 0.6mL DPBS in a 1mL open-top thickwall polycarbonate tube (Beckman Coulter, USA) and ultracentrifugation at 100,000 x g for 90 min at 4°C using an MLA-130 fixed angle rotor (Beckman Coulter, USA). The final EV pellet was resuspended in 100 μL of DPBS.

#### Ultrafiltration and size-exclusion chromatography (SEC)

Large volumes (>50 mL) of CCM were concentrated using Amicon® Ultra-15 Centrifugal Filters (100,000 MWCO) (Millipore Sigma, USA), according to manufacturer’s instructions. To minimize EV loss due to protein binding to the filter membranes, the latter were pre-treated with 5% Tween-80^107^ in DPBS and centrifuged at 2,608 x g (Eppendorf™ Centrifuge 5810 R, A-4-62 swinging bucket rotor) for 10 min at 4°C, followed by three washes in DPBS for 5 min at the same centrifugal speed. Once blocked, filter membranes were kept in DPBS until use, to prevent drying. The CCM was concentrated using these filters by centrifugation at 2,608 x g at 4°C. The time required to concentrate the material depended on the initial CCM volume, the final required retentate volume, the composition of the media being filtered. The concentrated CCM (retentate) was immediately transferred to a new tube, and the filter membrane was washed three times with DPBS to collect any loosely adherent material. EVs in the retentate were isolated by qEV Original 70nm size exclusion columns. SEC columns were stored in DPBS containing 0.05% sodium azide, at 4°C and used according to the manufacturer’s instructions. After column equilibration at r.t. and the initial flush of 10mL of DPBS, 0.5 mL of the retentate obtained by ultrafiltration was loaded onto the column and eluted with DPBS in 0.5 mL fractions. The presence of EVs in the various fractions was determined by measuring DiI (λ_Ex_/λ_Em_ = 540/580 nm) or DiD (λ_Ex_/λ_Em_ = 644/674 nm) fluorescence in each fraction. The protein content of each fraction was determined by measuring the absorbance at 280 nm using Nanodrop™ 2000c (Thermo Fisher Scientific, USA). Fractions enriched with EVs were pooled and concentrated using Amicon® Ultra-2mL Filters (10,000 MWCO) that were pre-treated with Tween-80 as described above. EV fractions were concentrated by centrifugation at 2,608 x g (with an A-4-62 swinging bucket rotor in an Eppendorf™ Centrifuge 5810 R) at 4°C.

### Storage and handling of EV samples

All EV samples were kept on ice and analyzed immediately after isolation for the determination of their number, size distribution and phenotype by imaging flow cytometry, ExoView, nanoparticle tracking analysis and dynamic light scattering. For other applications, EV samples were stored at -80°C until use. Repeated freeze-thaw cycles were avoided. To limit EV particle damage, EV samples were mixed by pipetting or by tube inversion, and never vortexed.

### Imaging Flow Cytometry EV Analysis (IFC)

IFC was used to determine EV number and detect mHTT-GFP within EVs. EV analysis by IFC was performed according to an established protocol^53^ using an Amnis® ImageStream mkII instrument, at the Flow Cytometry Core Facility of the Faculty of Medicine and Dentistry at the University of Alberta. All data were acquired with lasers set at maximum power, with 60x magnification, and the lowest flow rate. Data were acquired after 1 min of sample loading into the instrument to allow for flow stabilization. The DiI or DiD signal associated with EVs was acquired using a 560-595nm filter or a 642-745 nm filter, respectively. A 435-480 nm filter was used for brightfield imaging and a 745-780 nm filter was used for SSC measurements. GFP fluorescence in EVs carrying mHTT-GFP was detected using a 480-560 nm filter. DPBS pH 7.4 (BioSure® Sheath Solution, BioSure, UK) was used as sheath fluid. If required, samples were diluted with DPBS to avoid coincidence events. For most experiments, 3-10 technical replicates were acquired for each sample and data were averaged. In each experiment, 10, 000 events were acquired for the control sample, gating for events with a side scatter lower than that of the SpeedBeads (Amnis® SpeedBead® Kit for ImageStream®, Luminex Corporation, USA). All other samples were acquired for the same duration as the control. To confirm that the particles detected were membrane-bound EVs, each sample was re-analyzed after incubation with 0.05% NP-40 (IGEPAL® CA-630, filtered through 0.1μm PVDF filter) for 30 min at r.t., to solubilize all membrane-bound particles. Data were analysed using IDEAS® software v6.2. Any speed beads that may have been captured during the acquisition were gated out and masks were made to identify positive particles as described below:

- DiI: Intensity(M03,Ch03_DiI, 50-4095)
- DiD: Intensity(M11,Ch11_DiD, 60-4095)

As DiI or DiD were the fluorescent pan-EV markers, the spot count feature was added to their analysis to detect single particles.

### ExoView EV capture and analysis

Diluted cleared conditioned medium was incubated at r.t. overnight on ExoView microchips carrying immobilized antibodies against the tetraspanins CD9, CD63 and CD81. The next day, chips were washed using the ExoView C100 plate washer and incubated with fluorescent anti-tetraspanin antibodies. The EVs captured on the chip were sized by single particle interferometric reflectance imaging (SP-IRIS) and quantified using an ExoView R200 instrument. Data were analyzed using ExoView Analyzer 3.2.1.

### Detection and quantitation of ganglioside GM1

Relative quantification of GM1 in cell lysates and EV samples was performed by dot-blotting. Cells lysed in NP-40 lysis buffer (20 mM Tris-HCl at pH 7.4, 1% NP-40 v/v, 1 mM EDTA, 1 mM EGTA, 50 μM MG132, 1X protease/phosphatase inhibitor cocktail) were passed through a 27-G needle ten times, incubated on ice for 30 min and then sonicated at intensity 2.0 for 10 s using a Sonic Dismembrator Model 100 (Fisher Scientific, USA), prior to sample immobilization onto a 0.45μm pore size nitrocellulose membrane (Millipore, USA) using a 96-well Bio-Dot® apparatus (Bio-Rad, USA). EV samples were either sonicated at intensity 2.0 for 10 s or lysed with 0.01% NP-40. One microgram of protein from cell lysates was loaded in each well, in triplicates. EV samples were loaded in duplicates or triplicates. Membranes were blocked for 1 h with Odyssey® blocking buffer or Intercept® blocking buffer (LiCor Biosciences, USA), and incubated overnight at 4°C or for 2 h at r.t. with 0.25 μg/mL biotinylated cholera toxin B (Thermo Scientific, USA) diluted in Odyssey® or Intercept® blocking buffer to detect GM1, and with anti-α-tubulin antibodies (2125, Cell Signaling, USA) to control for equal loading of cell lysates. Membranes were then washed with TBS-T (20 mM Tris-HCl, 137 mM NaCl, pH 7.4-7.6 containing 0.1% Tween-20) and incubated with IRDye800®-conjugated Streptavidin (LiCor Biosciences, USA) diluted 1:5,000, and the appropriate fluorescent secondary antibody diluted 1:10,000, with Odyssey® or Intercept® blocking buffer:TBS-T (1:3) for 1 hour at r.t. Finally, membranes were washed once with TBS-T and once more with TBS before scanning in an Odyssey® near-infrared scanner (LiCor Biosciences, USA). Signal was quantified using Odyssey® Application Software v3.0 (LiCor Biosciences, USA).

### Statistical analysis

All statistical analyses were performed in GraphPad Prism v10. Unless otherwise stated, two-tailed paired *t*-test or ratio paired *t*-test were used for pair-wise comparisons. One-way ANOVA with Tukey’s post-hoc test, or repeated measures one-way ANOVA with Holm-Šidák post-hoc test were used to compare multiple groups. For comparing multiple groups with two or more independent variables, two-way ANOVA was used with Tukey’s *post-hoc* test.

**Additional materials and methods** are described in the Supplementary Information.

## Data availability

The data that support the findings of this study are available from the corresponding author upon reasonable request.

## Acknowledgements

This work was supported by grants from the Canadian Institutes for Health Research (CIHR), the Brain Canada Foundation, GlycoNET, the Alberta Prion Research Institute, and SynAD. JM was supported by a CIHR Banting and Best Canada Graduate Scholarship and by a FoMD Dean’s Doctoral Student Award. Flow cytometry and ExoView analyses were performed at the Flow Cytometry Facility of the Faculty of Medicine & Dentistry (FoMD) at the University of Alberta, which receives financial support from the FoMD and Canada Foundation for Innovation (CFI) awards to contributing investigators. NTA experiments were conducted at Nanostics Precision Health (Edmonton, Canada). We thank Dr. Aja Rieger for technical support and guidance for the analysis of EVs by IFC, Angelle Britton for training support in human iPSC culturing, and TRB Chemedica Int. for providing ganglioside GM1. Figures 2b, 5e and 6a were created, in part, with BioRender.

## Author information

JM, VK, LCM, JI, AKZ, DP and EM designed and performed experiments and analyzed data; DO, JH and AM performed experiments, VS and SAM provided expertise and supervised experiments; SS and EPC conceived research and designed experiments; SS supervised experiments and data analysis; SS and JM wrote the manuscript; all authors read and approved the final manuscript.

## Competing interests

SS holds a patent licensed to Zulia Inc. (US) for the use of GM1 for the treatment of Huntington’s disease. JM, DO and SS have filed a patent application for the use of GA1 for the treatment of breast cancer and other malignancies.

## Notes

### Competing Interest Statement

SS holds a patent licensed to Zulia Inc. (US) for the use of GM1 for the treatment of Huntington disease. JM, DO and SS have filed a patent application for the use of GA1 for the treatment of breast cancer and other malignancies.

## References

1 van Niel, G., D’Angelo, G. & Raposo, G. Shedding light on the cell biology of extracellular vesicles. Nature reviews. Molecular cell biology 19, 213–228 (2018). 10.1038/nrm.2017.125

2 Chang, W. H., Cerione, R. A. & Antonyak, M. A. Extracellular Vesicles and Their Roles in Cancer Progression. Methods Mol Biol 2174, 143–170 (2021). 10.1007/978-1-0716-0759-6_10

3 Takeuchi, T. Pathogenic and protective roles of extracellular vesicles in neurodegenerative diseases. J Biochem 169, 181–186 (2021). 10.1093/jb/mvaa131

4 Liang, W. et al. Mitochondria are secreted in extracellular vesicles when lysosomal function is impaired. Nat Commun 14, 5031 (2023). 10.1038/s41467-023-40680-5

5 Budnik, V., Ruiz-Canada, C. & Wendler, F. Extracellular vesicles round off communication in the nervous system. Nat Rev Neurosci 17, 160–172 (2016). 10.1038/nrn.2015.29

6 Bahram Sangani, N., Gomes, A. R., Curfs, L. M. G. & Reutelingsperger, C. P. The role of Extracellular Vesicles during CNS development. Prog Neurobiol 205, 102124 (2021). 10.1016/j.pneurobio.2021.102124

7 Bahrini, I., Song, J. H., Diez, D. & Hanayama, R. Neuronal exosomes facilitate synaptic pruning by up-regulating complement factors in microglia. Scientific reports 5, 7989 (2015). 10.1038/srep07989

8 Iguchi, Y. et al. Exosome secretion is a key pathway for clearance of pathological TDP-43. Brain 139, 3187–3201 (2016). 10.1093/brain/aww237

9 Kanemoto, S. et al. Multivesicular body formation enhancement and exosome release during endoplasmic reticulum stress. Biochem Biophys Res Commun 480, 166–172 (2016). 10.1016/j.bbrc.2016.10.019

10 Liu, S. et al. Highly efficient intercellular spreading of protein misfolding mediated by viral ligand-receptor interactions. Nat Commun 12, 5739 (2021). 10.1038/s41467-021-25855-2

11 Fevrier, B. et al. Cells release prions in association with exosomes. Proceedings of the National Academy of Sciences of the United States of America 101, 9683–9688 (2004). 10.1073/pnas.0308413101

12 Wang, Y. et al. The release and trans-synaptic transmission of Tau via exosomes. Molecular neurodegeneration 12, 5 (2017). 10.1186/s13024-016-0143-y

13 Sardar Sinha, M., et al. Alzheimer’s disease pathology propagation by exosomes containing toxic amyloid-beta oligomers. Acta Neuropathol 136, 41–56 (2018). 10.1007/s00401-018-1868-1

14 Wang, J. K. T., Langfelder, P., Horvath, S. & Palazzolo, M. J. Exosomes and Homeostatic Synaptic Plasticity Are Linked to Each other and to Huntington’s, Parkinson’s, and Other Neurodegenerative Diseases by Database-Enabled Analyses of Comprehensively Curated Datasets. Front Neurosci 11, 149 (2017). 10.3389/fnins.2017.00149

15 Estes, R. E., Lin, B., Khera, A. & Davis, M. Y. Lipid Metabolism Influence on Neurodegenerative Disease Progression: Is the Vehicle as Important as the Cargo? Frontiers in molecular neuroscience 14, 788695 (2021). 10.3389/fnmol.2021.788695

16 Hansen, S. B. & Wang, H. The shared role of cholesterol in neuronal and peripheral inflammation. Pharmacol Ther 249, 108486 (2023). 10.1016/j.pharmthera.2023.108486

17 Olesova, D. et al. Changes in lipid metabolism track with the progression of neurofibrillary pathology in tauopathies. J Neuroinflammation 21, 78 (2024). 10.1186/s12974-024-03060-4

18 Hallett, P. J., Engelender, S. & Isacson, O. Lipid and immune abnormalities causing age-dependent neurodegeneration and Parkinson’s disease. Journal of neuroinflammation 16, 153 (2019). 10.1186/s12974-019-1532-2

19 Record, M., Silvente-Poirot, S., Poirot, M. & Wakelam, M. J. O. Extracellular vesicles: lipids as key components of their biogenesis and functions. J Lipid Res 59, 1316–1324 (2018). 10.1194/jlr.E086173

20 Abdullah, M. et al. Cholesterol Regulates Exosome Release in Cultured Astrocytes. Front Immunol 12, 722581 (2021). 10.3389/fimmu.2021.722581

21 Jin, X. et al. Exosomal lipid PI4P regulates small extracellular vesicle secretion by modulating intraluminal vesicle formation. J Extracell Vesicles 12, e12319 (2023). 10.1002/jev2.12319

22 Beer, K. B. et al. Extracellular vesicle budding is inhibited by redundant regulators of TAT-5 flippase localization and phospholipid asymmetry. Proceedings of the National Academy of Sciences of the United States of America 115, E1127–E1136 (2018). 10.1073/pnas.1714085115

23 Dubois, L., Ronquist, K. K., Ek, B., Ronquist, G. & Larsson, A. Proteomic Profiling of Detergent Resistant Membranes (Lipid Rafts) of Prostasomes. Mol Cell Proteomics 14, 3015–3022 (2015). 10.1074/mcp.M114.047530

24 Salzer, U. & Prohaska, R. Stomatin, flotillin-1, and flotillin-2 are major integral proteins of erythrocyte lipid rafts. Blood 97, 1141–1143 (2001). 10.1182/blood.v97.4.1141

25 Fuki, I. V., Meyer, M. E. & Williams, K. J. Transmembrane and cytoplasmic domains of syndecan mediate a multi-step endocytic pathway involving detergent-insoluble membrane rafts. Biochem J 351 **Pt** **3**, 607–612 (2000).

26 Hemler, M. E., Tetraspanin functions and associated microdomains. Nature reviews. Molecular cell biology 6, 801–811 (2005). 10.1038/nrm1736

27 de Gassart, A., Geminard, C., Fevrier, B., Raposo, G. & Vidal, M. Lipid raft-associated protein sorting in exosomes. Blood 102, 4336–4344 (2003). 10.1182/blood-2003-03-0871

28 Pfrieger, F. W. & Vitale, N. Cholesterol and the journey of extracellular vesicles. *Journal of lipid research* (2018). 10.1194/jlr.R084210

29 Sipione, S., Monyror, J., Galleguillos, D., Steinberg, N. & Kadam, V. Gangliosides in the Brain: Physiology, Pathophysiology and Therapeutic Applications. Frontiers in neuroscience 14, 572965 (2020). 10.3389/fnins.2020.572965

30 Schnaar, R. L. The Biology of Gangliosides. Adv Carbohydr Chem Biochem 76, 113–148 (2019). 10.1016/bs.accb.2018.09.002

31 He, X., Guan, F. & Lei, L. Structure and function of glycosphingolipids on small extracellular vesicles. Glycoconjugate journal 39, 197–205 (2022). 10.1007/s10719-022-10052-0

32 Skryabin, G. O., Komelkov, A. V., Savelyeva, E. E. & Tchevkina, E. M. Lipid Rafts in Exosome Biogenesis. Biochemistry (Mosc) 85, 177–191 (2020). 10.1134/S0006297920020054

33 Sonnino, S., Mauri, L., Chigorno, V. & Prinetti, A. Gangliosides as components of lipid membrane domains. Glycobiology 17, 1R–13R (2007). 10.1093/glycob/cwl052

34 Dasgupta, R., Miettinen, M. S., Fricke, N., Lipowsky, R. & Dimova, R. The glycolipid GM1 reshapes asymmetric biomembranes and giant vesicles by curvature generation. Proc Natl Acad Sci U S A 115, 5756–5761 (2018). 10.1073/pnas.1722320115

35 Haraszti, R. A. et al. High-resolution proteomic and lipidomic analysis of exosomes and microvesicles from different cell sources. J Extracell Vesicles 5, 32570 (2016). 10.3402/jev.v5.32570

36 Desplats, P. A. et al. Glycolipid and ganglioside metabolism imbalances in Huntington’s disease. Neurobiol Dis 27, 265–277 (2007). S0969-9961(07)00107-6 [pii] 10.1016/j.nbd.2007.05.003

37 Huebecker, M. et al. Reduced sphingolipid hydrolase activities, substrate accumulation and ganglioside decline in Parkinson’s disease. Mol Neurodegener 14, 40 (2019). 10.1186/s13024-019-0339-z

38 Seyfried, T. N. et al. Sex-Related Abnormalities in Substantia Nigra Lipids in Parkinson’s Disease. ASN neuro 10, 1759091418781889 (2018). 10.1177/1759091418781889

39 Denny, C. A., Desplats, P. A., Thomas, E. A. & Seyfried, T. N. Cerebellar lipid differences between R6/1 transgenic mice and humans with Huntington’s disease. J Neurochem 115, 748–758 (2010). 10.1111/j.1471-4159.2010.06964.x

40 Maglione, V. et al. Impaired ganglioside metabolism in Huntington’s disease and neuroprotective role of GM1. J Neurosci 30, 4072–4080 (2010). 10.1523/jneurosci.6348-09.2010

41 Boukhris, A. et al. Alteration of ganglioside biosynthesis responsible for complex hereditary spastic paraplegia. Am J Hum Genet 93, 118–123 (2013). 10.1016/j.ajhg.2013.05.006

42 Harlalka, G. V. et al. Mutations in B4GALNT1 (GM2 synthase) underlie a new disorder of ganglioside biosynthesis. Brain 136, 3618–3624 (2013). 10.1093/brain/awt270

43 Boccuto, L. et al. A mutation in a ganglioside biosynthetic enzyme, ST3GAL5, results in salt & pepper syndrome, a neurocutaneous disorder with altered glycolipid and glycoprotein glycosylation. Hum Mol Genet 23, 418–433 (2014). 10.1093/hmg/ddt434

44 Simpson, M. A. et al. Infantile-onset symptomatic epilepsy syndrome caused by a homozygous loss-of-function mutation of GM3 synthase. Nat Genet 36, 1225–1229 (2004). http://ng1460

45 Alpaugh, M. et al. Disease-modifying effects of ganglioside GM1 in Huntington’s disease models. EMBO molecular medicine 9, 1537–1557 (2017). 10.15252/emmm.201707763

46 Di Pardo, A. et al. Ganglioside GM1 induces phosphorylation of mutant huntingtin and restores normal motor behavior in Huntington disease mice. Proc Natl Acad Sci U S A 109, 3528–3533 (2012). 10.1073/pnas.1114502109

47 Schneider, J. S. et al. GM1 Ganglioside Modifies alpha-Synuclein Toxicity and is Neuroprotective in a Rat alpha-Synuclein Model of Parkinson’s Disease. Scientific reports 9, 8362 (2019). 10.1038/s41598-019-42847-x

48 Schneider, J. S. et al. A randomized, controlled, delayed start trial of GM1 ganglioside in treated Parkinson’s disease patients. J Neurol Sci 324, 140–148 (2013). 10.1016/j.jns.2012.10.024

49 Itokazu, Y., Fuchigami, T., Morgan, J. C. & Yu, R. K. Intranasal infusion of GD3 and GM1 gangliosides downregulates alpha-synuclein and controls tyrosine hydroxylase gene in a PD model mouse. Mol Ther 29, 3059–3071 (2021). 10.1016/j.ymthe.2021.06.005

50 Leskawa, K. C., Erwin, R. E., Leon, A., Toffano, G. & Hogan, E. L. Incorporation of exogenous ganglioside GM1 into neuroblastoma membranes: inhibition by calcium ion and dependence upon membrane protein. Neurochem Res 14, 547–554 (1989). 10.1007/BF00964917

51 Lannigan, J. & Erdbruegger, U. Imaging flow cytometry for the characterization of extracellular vesicles. Methods 112, 55–67 (2017). 10.1016/j.ymeth.2016.09.018

52 Görgens, A. et al. Optimisation of imaging flow cytometry for the analysis of single extracellular vesicles by using fluorescence-tagged vesicles as biological reference material. J Extracell Vesicles 8, 1587567 (2019). 10.1080/20013078.2019.1587567

53 Viveiros, A. et al. In-Cell Labeling Coupled to Direct Analysis of Extracellular Vesicles in the Conditioned Medium to Study Extracellular Vesicles Secretion with Minimum Sample Processing and Particle Loss. Cells 11, 351 (2022). 10.3390/cells11030351

54 Monguio-Tortajada, M., Galvez-Monton, C., Bayes-Genis, A., Roura, S. & Borras, F. E. Extracellular vesicle isolation methods: rising impact of size-exclusion chromatography. Cell Mol Life Sci 76, 2369–2382 (2019). 10.1007/s00018-019-03071-y

55 Liangsupree, T., Multia, E. & Riekkola, M. L. Modern isolation and separation techniques for extracellular vesicles. J Chromatogr A 1636, 461773 (2021). 10.1016/j.chroma.2020.461773

56 Mathieu, M. et al. Specificities of exosome versus small ectosome secretion revealed by live intracellular tracking of CD63 and CD9. Nature communications 12, 4389 (2021). 10.1038/s41467-021-24384-2

57 Kowal, J. et al. Proteomic comparison defines novel markers to characterize heterogeneous populations of extracellular vesicle subtypes. Proceedings of the National Academy of Sciences of the United States of America 113, E968–977 (2016). 10.1073/pnas.1521230113

58 Gardiner, C., Ferreira, Y. J., Dragovic, R. A., Redman, C. W. & Sargent, I. L. Extracellular vesicle sizing and enumeration by nanoparticle tracking analysis. J Extracell Vesicles 2 (2013). 10.3402/jev.v2i0.19671

59 Khan, M. A., Anand, S., Deshmukh, S. K., Singh, S. & Singh, A. P. Determining the Size Distribution and Integrity of Extracellular Vesicles by Dynamic Light Scattering. Methods Mol Biol 2413, 165–175 (2022). 10.1007/978-1-0716-1896-7_17

60 Arab, T. et al. Characterization of extracellular vesicles and synthetic nanoparticles with four orthogonal single-particle analysis platforms. J Extracell Vesicles 10, e12079 (2021). 10.1002/jev2.12079

61 Breitwieser, K. et al. Detailed Characterization of Small Extracellular Vesicles from Different Cell Types Based on Tetraspanin Composition by ExoView R100 Platform. Int J Mol Sci 23 (2022). 10.3390/ijms23158544

62 Daaboul, G. G. et al. Digital Detection of Exosomes by Interferometric Imaging. Scientific reports 6, 37246 (2016). 10.1038/srep37246

63 Mathieu, M. et al. Specificities of exosome versus small ectosome secretion revealed by live intracellular tracking of CD63 and CD9. Nature Communications 12, 4389 (2021). 10.1038/s41467-021-24384-2

64 Kowal, J. et al. Proteomic comparison defines novel markers to characterize heterogeneous populations of extracellular vesicle subtypes. Proceedings of the National Academy of Sciences 113, E968–E977 (2016). 10.1073/pnas.1521230113

65 Zhao, H. et al. Inhibiting glycosphingolipid synthesis improves glycemic control and insulin sensitivity in animal models of type 2 diabetes. Diabetes 56, 1210–1218 (2007). 10.2337/db06-0719

66 Di Pardo, A., Amico, E. & Maglione, V. Impaired Levels of Gangliosides in the Corpus Callosum of Huntington Disease Animal Models. Frontiers in neuroscience 10, 457 (2016). 10.3389/fnins.2016.00457

67 Segler-Stahl, K., Webster, J. C. & Brunngraber, E. G. Changes in the concentration and composition of human brain gangliosides with aging. Gerontology 29, 161–168 (1983).

68 Wu, G., Lu, Z. H., Kulkarni, N. & Ledeen, R. W. Deficiency of ganglioside GM1 correlates with Parkinson’s disease in mice and humans. J Neurosci Res 90, 1997–2008 (2012). 10.1002/jnr.23090

69 Zaprianova, E. et al. Serum ganglioside patterns in multiple sclerosis. Neurochem Res 26, 95–100 (2001).

70 Schmidt, E. N. et al. Siglec-6 mediates the uptake of extracellular vesicles through a noncanonical glycolipid binding pocket. Nature communications 14, 2327 (2023). 10.1038/s41467-023-38030-6

71 Deng, J. et al. Neurons Export Extracellular Vesicles Enriched in Cysteine String Protein and Misfolded Protein Cargo. Scientific reports 7, 956 (2017). 10.1038/s41598-017-01115-6

72 Apostol, B. L. et al. A cell-based assay for aggregation inhibitors as therapeutics of polyglutamine-repeat disease and validation in Drosophila. Proceedings of the National Academy of Sciences of the United States of America 100, 5950–5955 (2003). 10.1073/pnas.2628045100

73 Trettel, F. et al. Dominant phenotypes produced by the HD mutation in STHdh(Q111) striatal cells. Hum Mol Genet 9, 2799–2809 (2000).

74 Ledeen, R. W., Kopitz, J., Abad-Rodriguez, J. & Gabius, H. J. Glycan Chains of Gangliosides: Functional Ligands for Tissue Lectins (Siglecs/Galectins). Prog Mol Biol Transl Sci 156, 289–324 (2018). 10.1016/bs.pmbts.2017.12.004

75 Royo, F., Théry, C., Falcón-Pérez, J. M., Nieuwland, R. & Witwer, K. W. Methods for Separation and Characterization of Extracellular Vesicles: Results of a Worldwide Survey Performed by the ISEV Rigor and Standardization Subcommittee. Cells 9 (2020). 10.3390/cells9091955

76 Ma, Q. et al. Vesicular Ganglioside GM1 From Breast Tumor Cells Stimulated Epithelial-to-Mesenchymal Transition of Recipient MCF-10A Cells. Frontiers in oncology 12, 837930 (2022). 10.3389/fonc.2022.837930

77 Sung, B. H. et al. A live cell reporter of exosome secretion and uptake reveals pathfinding behavior of migrating cells. Nat Commun 11, 2092 (2020). 10.1038/s41467-020-15747-2

78 Chinnapen, D. J. et al. Lipid sorting by ceramide structure from plasma membrane to ER for the cholera toxin receptor ganglioside GM1. Dev Cell 23, 573–586 (2012). 10.1016/j.devcel.2012.08.002

79 Trajkovic, K. et al. Ceramide triggers budding of exosome vesicles into multivesicular endosomes. Science 319, 1244–1247 (2008). 10.1126/science.1153124

80 Sandhoff, R., Schulze, H. & Sandhoff, K. Ganglioside Metabolism in Health and Disease. Progress in molecular biology and translational science 156, 1–62 (2018). 10.1016/bs.pmbts.2018.01.002

81 Verderio, C., Gabrielli, M. & Giussani, P. Role of sphingolipids in the biogenesis and biological activity of extracellular vesicles. J Lipid Res 59, 1325–1340 (2018). 10.1194/jlr.R083915

82 Rodi, P. M., Maggio, B. & Bagatolli, L. A. Direct visualization of the lateral structure of giant vesicles composed of pseudo-binary mixtures of sulfatide, asialo-GM1 and GM1 with POPC. Biochim Biophys Acta Biomembr 1860, 544–555 (2018). 10.1016/j.bbamem.2017.10.022

83 Carey, D. J., Bendt, K. M. & Stahl, R. C. The cytoplasmic domain of syndecan-1 is required for cytoskeleton association but not detergent insolubility. Identification of essential cytoplasmic domain residues. J Biol Chem 271, 15253–15260 (1996). 10.1074/jbc.271.25.15253

84 Zhang, G. L. et al. The Human Ganglioside Interactome in Live Cells Revealed Using Clickable Photoaffinity Ganglioside Probes. Journal of the American Chemical Society 146, 17801–17816 (2024). 10.1021/jacs.4c03196

85 Mitsuda, T. et al. Overexpression of ganglioside GM1 results in the dispersion of platelet-derived growth factor receptor from glycolipid-enriched microdomains and in the suppression of cell growth signals. J Biol Chem 277, 11239–11246 (2002). 10.1074/jbc.M107756200

86 Prendergast, J. et al. Ganglioside regulation of AMPA receptor trafficking. The Journal of neuroscience : the official journal of the Society for Neuroscience 34, 13246–13258 (2014). 10.1523/JNEUROSCI.1149-14.2014

87 Julien, S., Bobowski, M., Steenackers, A., Le Bourhis, X. & Delannoy, P. How Do Gangliosides Regulate RTKs Signaling? Cells 2, 751–767 (2013). 10.3390/cells2040751

88 Kabayama, K. et al. Dissociation of the insulin receptor and caveolin-1 complex by ganglioside GM3 in the state of insulin resistance. Proceedings of the National Academy of Sciences of the United States of America 104, 13678–13683 (2007). 10.1073/pnas.0703650104

89 Posse de Chaves, E. & Sipione, S. Sphingolipids and gangliosides of the nervous system in membrane function and dysfunction. FEBS Lett 584, 1748–1759 (2010). 10.1016/j.febslet.2009.12.010

90 Kracun, I. et al. Gangliosides in the human brain development and aging. Neurochem Int 20, 421–431 (1992).

91 Hong, Y., Zhao, T., Li, X. J. & Li, S. Mutant Huntingtin Inhibits alphaB-Crystallin Expression and Impairs Exosome Secretion from Astrocytes. The Journal of neuroscience : the official journal of the Society for Neuroscience 37, 9550–9563 (2017). 10.1523/JNEUROSCI.1418-17.2017

92 Beatriz, M. et al. Defective mitochondria-lysosomal axis enhances the release of extracellular vesicles containing mitochondrial DNA and proteins in Huntington’s disease. Journal of Extracellular Biology 1, e65 (2022).

93 Takov, K., Yellon, D. M. & Davidson, S. M. Comparison of small extracellular vesicles isolated from plasma by ultracentrifugation or size-exclusion chromatography: yield, purity and functional potential. J Extracell Vesicles 8, 1560809 (2019). 10.1080/20013078.2018.1560809

94 Baixauli, F., Lopez-Otin, C. & Mittelbrunn, M. Exosomes and autophagy: coordinated mechanisms for the maintenance of cellular fitness. Frontiers in immunology 5, 403 (2014). 10.3389/fimmu.2014.00403

95 Ledeen, R. W. & Wu, G. The multi-tasked life of GM1 ganglioside, a true factotum of nature. Trends Biochem Sci 40, 407–418 (2015). 10.1016/j.tibs.2015.04.005

96 Pink, D., Donnelier, J., Lewis, J. D. & Braun, J. E. A. Cysteine String Protein Controls Two Routes of Export for Misfolded Huntingtin. Frontiers in neuroscience 15, 762439 (2021). 10.3389/fnins.2021.762439

97 Isogai, T. et al. Extracellular vesicles adhere to cells predominantly through the interaction of CD151-associated integrin heterodimers and GM1 with laminin. *bioRxiv*, 2024.2004.2011.589011 (2024). 10.1101/2024.04.11.589011

98 Maacha, S. et al. Extracellular vesicles-mediated intercellular communication: roles in the tumor microenvironment and anti-cancer drug resistance. Mol Cancer 18, 55 (2019). 10.1186/s12943-019-0965-7

99 Webber, J., Yeung, V. & Clayton, A. Extracellular vesicles as modulators of the cancer microenvironment. Semin Cell Dev Biol 40, 27–34 (2015). 10.1016/j.semcdb.2015.01.013

100 Toyoshima, M. et al. Inhibition of tumor growth and metastasis by depletion of vesicular sorting protein Hrs: its regulatory role on E-cadherin and beta-catenin. Cancer Res 67, 5162–5171 (2007). 10.1158/0008-5472.CAN-06-2756

101 Koo, T. H. et al. Syntenin is overexpressed and promotes cell migration in metastatic human breast and gastric cancer cell lines. Oncogene 21, 4080–4088 (2002). 10.1038/sj.onc.1205514

102 Brewer, G. J., Torricelli, J. R., Evege, E. K. & Price, P. J. Optimized survival of hippocampal neurons in B27-supplemented Neurobasal, a new serum-free medium combination. Journal of neuroscience research 35, 567–576 (1993). 10.1002/jnr.490350513

103 Sciannamblo, M. et al. Changes of the ganglioside pattern and content in human fibroblasts by high density cell population subculture progression. Glycoconjugate journal 19, 181–186 (2002). 10.1023/A:1024249707516

104 Wang, C. et al. Scalable Production of iPSC-Derived Human Neurons to Identify Tau-Lowering Compounds by High-Content Screening. Stem Cell Reports 9, 1221–1233 (2017). 10.1016/j.stemcr.2017.08.019

105 Kang, S. G. et al. Pathologic tau conformer ensembles induce dynamic, liquid-liquid phase separation events at the nuclear envelope. BMC Biol 19, 199 (2021). 10.1186/s12915-021-01132-y

106 Thery, C., Amigorena, S., Raposo, G. & Clayton, A. Isolation and characterization of exosomes from cell culture supernatants and biological fluids. *Current protocols in cell biology* **Chapter 3**, Unit 3 22 (2006). 10.1002/0471143030.cb0322s30

107 Lee, K. J. et al. Modulation of nonspecific binding in ultrafiltration protein binding studies. Pharmaceutical research 20, 1015–1021 (2003). 10.1023/a:1024406221962

