## Supplementary methods, table and figures for "Gangliosides Modulate the Secretion of Extracellular Vesicles and Their Misfolded Protein Cargo"

### Chemicals, reagents, and materials

Purified porcine GM1 was provided by TRB Chemedica Int., Switzerland. Purified bovine GM3 and GD3 were purchased from Avanti Polar Lipids Inc., USA. Purified bovine GM2, GD2 and GQ1b were purchased from Cayman Chemical, USA. Purified GD1a, GD1b and GT1b, semisynthetic asialo-GM1 and GM1 pentasaccharide were purchased from Enzo Life Sciences, USA. Genz-123346 was purchased from Toronto Research Chemicals, Canada. Vybrant™ DiI and Vybrant™ DiD Cell-Labeling solutions, mouse laminin, and DMEM/F12-HEPES (phenol red free) were purchased from Thermo Fisher Scientific, USA. L-Glutamine, sodium pyruvate, Geneticin®, N-2 supplement, B27 supplement, GlutaMAX™ supplement, Opti-MEM® I reduced serum media (with phenol red), Opti-MEM® I reduced serum media (phenol red-free), Dulbecco's Modified Eagle Medium (DMEM) – high glucose (phenol red-free), Minimum Essential Medium (MEM) (with phenol red), MEM (phenol red-free), MEM Non-Essential Amino Acids Solution, Hank's Balanced Salt Solution (HBSS), Neurobasal™-A medium (phenol red-free and with phenol red), Phosphate Buffered Saline (without calcium and magnesium), MEM Non-Essential Amino Acids Solution (NEAA), Essential 8™ Medium, KnockOut™ DMEM, StemPro Accutase, and horse serum were purchased from Gibco™, Thermo Fisher Scientific, USA. Cell-culture grade DMSO, Tween-80, fetal bovine serum (FBS), EDTA, EGTA, MG132, bovine serum albumin (BSA), BSA free of fatty acids, and Amicon® Ultra-15 Centrifugal Filters (10,000 MWCO or 100,000 MWCO) were purchased from Sigma Aldrich, USA. Doxycycline hyclate was purchased from either Gibco™, Thermo Fisher Scientific, USA (for Tau-inducible cells) or Sigma Aldrich, USA (for iPSCs). Recombinant Human Neurotrophin-3 (NT-3), Recombinant Human brain derived neurotrophic factor (BDNF) were purchased from Peprotech. Y-27632 dihydrochloride was purchased from MedChemExpress. DMEM – high glucose (with phenol red), cell-culture grade Dulbecco's modified phosphate buffered saline (DPBS), Penicillin/Streptomycin were purchased from HyClone™, GE Healthcare, USA. Complete® Protease inhibitor cocktail and PhosStop® phosphatase inhibitor cocktail were purchased from Roche, Switzerland. IGEPAL® CA-630 was from Sigma Aldrich, USA. Nitrocellulose membranes were purchased from Bio-Rad, USA, while PVDF membranes (Immobilon®-FL) were from Merck Millipore, USA. Pre-lubricated microcentrifuge tubes (CoStar, Corning Life Sciences, USA) were used for the isolation and analysis of EVs. All polystyrene and polypropylene tubes were from Falcon™, Thermo Fisher Scientific, USA. qEV Original 70 nm columns were purchased from IZON Science, Ltd., USA. ExoView human tetraspanin plasma Kits were purchased from Unchained Laboratories, USA.

**Fluorometry**

DiI or DiD fluorescence was measured in cell lysates and EV fractions using a SpectraMax® i3x multi-mode microplate reader (Molecular Devices, USA). Ex/Em for DiI is 540/580nm while that for DiD is 644/674nm. The bandwidth of all excitation and emission wavelengths was set to 15nm. Fluorescence was measured in 96-well black-bottom plates by well-scan reading.

**CytoFlex EV Analysis**

Unlabeled EV samples (SEC-isolated or CCM) were analyzed using the CytoFLEX S (Beckman Coulter, USA) instrument at the Flow Cytometry Core Facility of the Faculty of Medicine and Dentistry at the University of Alberta. Samples were diluted 1/500 (SEC-isolated) or 1/10 (CCM) with 0.1 μm-filtered DPBS. Following flow stabilization, each sample was recorded for 60 seconds on slow speed and triggered with the 405 nm violet side scatter (V-SSC) laser with a detection threshold of 1750. The V-SSC gain was set to 200. Data analysis was performed using CytExpert software. V-SSC$^+$ events that were resolved from the electronic noise were gated based on DPBS for SEC-isolated samples or unconditioned collection medium diluted 1/10 with DPBS for CCM samples.

**Nanoparticle Tracking Analysis (NTA)**

EV number and size distribution were determined using the NanoSight LM10 system (405nm laser, software version 3.3 – Sample Assistant Dev Build 3.3.203) (NanoSight, United Kingdom). Instrument sizing was checked using 200 nm polystyrene beads (Nanosphere, USA) resuspended in 10 mM potassium chloride (KCl). CCM samples were diluted with 0.1 µm-filtered PBS (Gibco™, Thermo Fisher Scientific, USA) to acquire 20-90 particles/frame. 300 µL of the sample was injected into the NanoSight chamber with a 1mL syringe. Five serial measurements were taken for each sample, and each measurement was taken for 1 min. PBS was injected into the chamber to wash the system in between samples.

**Dynamic Light Scattering (DLS)**

Freshly prepared retentate obtained by ultrafiltration of the CCM was used for DLS measurements to determine EV size distribution. An aliquot of the retentate corresponding to 350 µg of proteins (measured by BCA protein assay) was subjected to asymmetric flow field-flow fractionation on an AF2000 Postnova system, using DPBS, pH 7.4, as running buffer. Samples were focussed for 5 min before elution at a flow rate of 0.5 mL/min. A slot pump was run at 0.3 mL/min to concentrate the samples before passing them through the detectors. Fractions of 0.2 mL were collected and analyzed by an in-line DAWN HELEOS II detector (Wyatt Technology) and using ASTRA software v6.1.

### Protein quantification

Protein concentration in cell lysates and EV samples was measured with a Pierce™ BCA Protein Assay or with a Pierce™ Enhanced BCA Protein Assay (Thermo Fisher Scientific, USA), respectively, according to manufacturer's instructions. All samples for protein measurements were loaded in triplicates. The absorbance of each sample was measured by endpoint reading at 562nm, using a SpectraMax® i3x multi-mode microplate reader (Molecular Devices, USA).

### Immunoblotting

Cell lysates (30-60µg of proteins) and equal fractions of EV samples isolated by UC or by SEC (not exceeding 60 µg of total EV proteins) were denatured in sample loading buffer (62.5 mM Tris-HCl pH 6.8, 2% SDS, 10% glycerol, 5% 2-mercaptoethanol, 6.25 mg/mL bromophenol blue) for 5 min at 95°C. Samples were loaded onto 4-20% SDS PAGE gradient gels and run in a Mini-PROTEAN® II electrophoresis chamber (Bio-Rad, USA). Proteins were transferred to a 0.45µm PVDF membrane in Towbin buffer (25 mM Tris-HCl, 192 mM glycine, 20% methanol v/v) using a Mini Trans-Blot® apparatus (Bio-Rad, USA). For full-length huntingtin immunoblots, samples were loaded onto 4-12% SDS PAGE gels and run as above, and protein transfer was performed using a modified Towbin buffer (25 mM Tris-HCl, 192mM glycine, 16% methanol v/v, 0.5% SDS). PVDF membranes were blocked for 1 h with 5% BSA in TBS-T, Odyssey®, or Intercept® blocking buffer (LiCor Biosciences, USA), and then incubated with primary antibodies for 2 h at r.t. or overnight at 4°C. The complete list of antibodies used can be found in the Supplementary Table 1. The membranes were then washed with TBS-T and incubated with the appropriate fluorescent secondary antibodies (LiCor BioSciences, USA) diluted in blocking buffer:TBS-T (1:3) for 1 h at r.t. After washing with TBS-T and TBS, the membranes were scanned in an

Odyssey® or Odyssey® M near-infrared scanner (LiCor Biosciences, USA) and signal was quantified using Odyssey® Application Software v3.0 (LiCor Biosciences, USA), Empiria Studio v3.1 or using ImageJ® software (NIH, USA). Signal from cell lysates was normalized over Revert™ total protein stain (LiCor Biosciences, USA) or tubulin signal as indicated.

**ELISA for EGFP-conjugated proteins**

Levels of mHTT-EGFP, WT or mutant tau-EGFP or A53T α-synuclein-EGFP in EVs isolated from the CCM by SEC were measured by the GFP SimpleStep ELISA® kit (Abcam, USA), following manufacturer's instructions. All materials and reagents were equilibrated to r.t. before use. The endpoint reading was recorded by measuring absorbance at 450 nm using SpectraMax® i3x multi-mode microplate reader (Molecular Devices, USA). Data were normalized over total cellular protein content.

**Proteinase K protection assay**

Two million N2a 97Q cells were seeded in each of twelve 100 mm dishes. After 2 h to allow for cell attachment, the cells were incubated with GM1 (50 µM) in SFM+N-2 for 6 h or left untreated (control). GM1 was washed off and EVs were collected in the conditioned medium for 16 h. and isolated by SEC as described above. Proteinase K (Sigma Aldrich, USA) was solubilized in 25 mM Tris-HCL, pH 8.0, containing 5 mM $CaCl_2$ (Buffer A) to a concentration of 0.1 mg/mL. For each experimental group, three EV aliquots (20 µg each) were freshly prepared and treated as follows. Aliquot 1: incubation with 2.0 µg of proteinase K in Buffer A for 1 h at 37°C, to digest accessible proteins on the EV surface, followed by enzyme inactivation at 95°C for 10 min and sample incubation with 0.1% saponin (Sigma Aldrich, USA) for 20 min at 37°C to permeabilize EV membranes and allow for the detection of EV luminal proteins. Aliquot 2: incubation with

0.1% saponin for 20 min at 37°C to permeabilize EV membranes and allow access to the luminal content, followed by treatment with 2.0 μg of proteinase K and heat inactivation. This results in the digestion of all EV proteins (luminal and membrane associated). Aliquot 3: Incubation with 0.1% saponin for 20 min at 37ºC to permeabilize EV membranes, followed by incubation in buffer A (equivalent volume to the proteinase K-treated samples) for 1h at 37ºC and heating at 95°C for 10 min.

Following treatment, NP-40 was added to all samples at a final concentration of 0.05% and incubated for 30 min at r.t. to lyse EVs completely. Samples were spotted on a 0.45 μm pore size nitrocellulose membrane (Millipore, USA) using a 96-well Bio-Dot® apparatus (Bio-Rad, USA), after dilution with DPBS to reduce the final concentration of NP-40 and saponin to no higher than 0.01% and 0.0167%, respectively, to avoid detergent interference with sample binding to the membrane. The membrane was blocked for 1 h with Intercept® blocking buffer (LiCor Biosciences, USA) and incubated with anti-Alix (611621, BD Biosciences, USA) and anti-HTT N17 antibodies (a gift from Dr. Ray Truant, McMaster University, Canada) overnight at 4°C. The membrane was then washed with TBS-T and incubated with fluorescent secondary antibodies (LiCor Biosciences, USA) for 1 h at r.t. After washing again with TBS-T and TBS, the membrane was scanned in an Odyssey® near-infrared scanner (LiCor Biosciences, USA). Signal was quantified using ImageJ® software (NIH, USA).

### Supplementary figure legends

**Supplementary figure 1. Imaging flow cytometry gating strategy for DiD- and DiI-stained EVs**

Gating strategy for DiD$^+$-EVs **(a)** DiI$^+$-EVs (**b**). Low side scatter events were gated to remove speedbeads ("R1" gate). An intensity mask was created to identify events with a fluorescent intensity above background, between 60 and 4095 for DiD$^+$-events or between 50 and 4095 for DiI$^+$-events. The spot count feature was used to identify and gate for single fluorescent events and quantify DiD$^+$- or DiI$^+$-EVs (R1 & single).

**Supplementary figure 2. Sizing of EVs from cells treated with GM1**

**a-b.** Representative histograms of the hydrodynamic radius distribution of EVs secreted by control and GM1-treated (50 μM) N2a cells (**a**) and control and GM1-treated N2a 97Q (**b**), as measured by dynamic light scattering (DLS). The tables show mean radius values ± SD for each group (n=1-2 independent experiments).

**c.** Particle size distribution profile of EVs secreted by control and GM1-treated N2a 97Q and measured by nanoparticle tracking analysis (NTA). EV concentration is normalized to total cellular protein content. The table shows the mean particle diameter ± SD and *p*-values obtained by two-tailed paired *t*-test (n=3 independent experiments).

**d.** Particle size distribution profile of EVs secreted by untreated and GM1-treated HD fibroblasts as measured by ExoView. EV counts are normalized to total cellular protein content. The table shows the mean particle diameter ± SD and *p*-values obtained by two-tailed paired *t*-test (n=3 biological replicates).

**Supplementary figure 3. Quantification of unlabelled EVs from Genz-123346-treated and Δ*B4galnt1* N2a cells**

Gangliosides were depleted from N2a cells by treatment with Genz-123356 or *B4galnt1* knockout. Unlabelled EVs were analyzed following SEC purification or directly from the CCM by micro-flow cytometry (CytoFLEX).

**a.** Representative histograms of violet side scatter (V-SSC) intensity of DPBS (left), SEC-isolated EVs from DMSO-treated (middle) and Genz-123346-treated N2a cells (right). The horizontal line shows the gate used to quantify EVs.

**b.** Quantification of the number of V-SSC$^+$-EVs secreted by DMSO- or Genz-123346-treated N2a cells normalized to total cellular protein content (n=5 technical replicates).

**c.** Representative histograms of V-SSC intensity of collection medium (left), CCM from WT N2a cells (middle), and Δ*B4galnt1* N2a cells (right).

**d.** Quantification of the number of V-SSC$^+$ EVs secreted by WT or Δ*B4galnt1* N2a cells normalized to total cellular protein content (n=5 technical replicates).

Bars indicate means ± SD.

**Supplementary figure 4. Tetraspanin profile of human fibroblast EVs detected by ExoView**

EVs were collected from 3 healthy and 3 HD fibroblast cell lines. Tetraspanins were profiled by ExoView.

**a.** Number of EVs bearing CD63, CD81 or CD9 normalized to total cellular protein content.

**b.** Number of EVs bearing only one tetraspanin, either CD63, CD81, or CD9, normalized to total cellular protein content.

**c.** Number of EVs bearing the indicated combinations of tetraspanins, normalized to total cellular protein content.

Bars are mean values ± SD. *$p$<0.05, by two-tailed unpaired *t*-test.

**Supplementary figure 5. The majority of mHTT associated with EVs is in the EV lumen**

EVs from N2a 97Q cells were treated with GM1 for 6 h and collected in phenol red-free, serum-free medium supplemented with N-2 for 16 h prior to isolation by SEC. EVs from control and GM1-treated cells were subjected to proteinase K digestion (0.02μg/μg of EVs) before or after EV membrane permeabilization with 0.1% saponin. In the absence of prior saponin permeabilization, proteinase K only digests proteins on the membrane and not in the lumen.

**a**. Representative dot blot showing the effects of proteinase K on mHTT and Alix (a luminal protein) in EVs treated as indicated. Alix was used as a control for proteinase K digestion. The blank is PBS, the EV resuspension buffer.

**b**. Densitometric analysis for mHTT (left) and Alix (right) in EVs, expressed as fold-change of undigested (heat + saponin group) untreated control (n=2 independent experiments for mHTT, 1 for Alix). Bars are means ± SD.

### Supplementary Table I - List of antibodies used

| Antibody | Host | Catalog no. | Source | Dilution | Application |
|---|---|---|---|---|---|
| **A1P1 (ALIX)** | Mouse | 611621 | BD Biosciences | 1:250 | Western Blotting and Proteinase K Protection Assay |
| **Calnexin** | Rabbit | ADI-SPA-860 | Enzo | 1:2000 | Western Blotting |
| **CD9** | Rabbit | ab92726 | Abcam | 1:2000 | Western Blotting |
| **Flotillin-1** | Mouse | 610820 | BD Biosciences | 1:1000 | Western Blotting |
| **GFP** | Rabbit | 2956 | Cell Signaling Technology | 1:2000 | Western Blotting |
| **GFP** | Mouse | sc-9996 | Santa Cruz | 1:500 | Western Blotting |
| **Huntingtin N17** | Rabbit | - | Dr. Truant, McMaster University | 1:5000-1:10000 | Western Blotting |
| **Huntingtin** | Mouse | MAB2166 | Millipore Sigma | 1:1000 | Western Blotting |
| **Tubulin** | Rabbit | 2125 | Cell Signaling Technology | 1:1000 | Western Blotting |
| **TSG101** | Rabbit | ab125011 | Abcam | 1:1000 | Western Blotting |
| **α-synuclein** | Rabbit | 2125 | Cell Signaling Technology | 1:10000 | Dot Blotting |

Supplementary figure 1

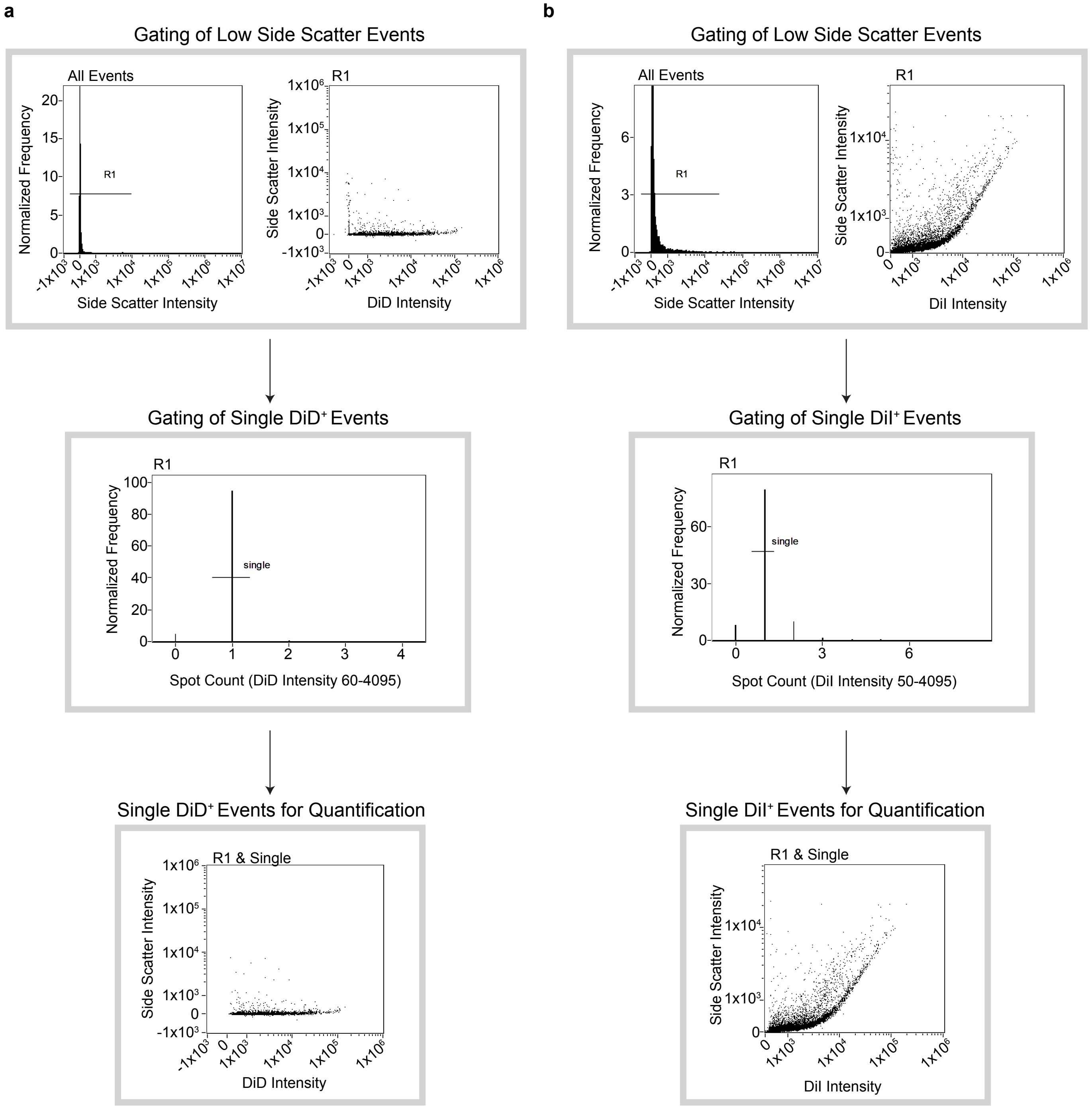

Supplementary figure 2

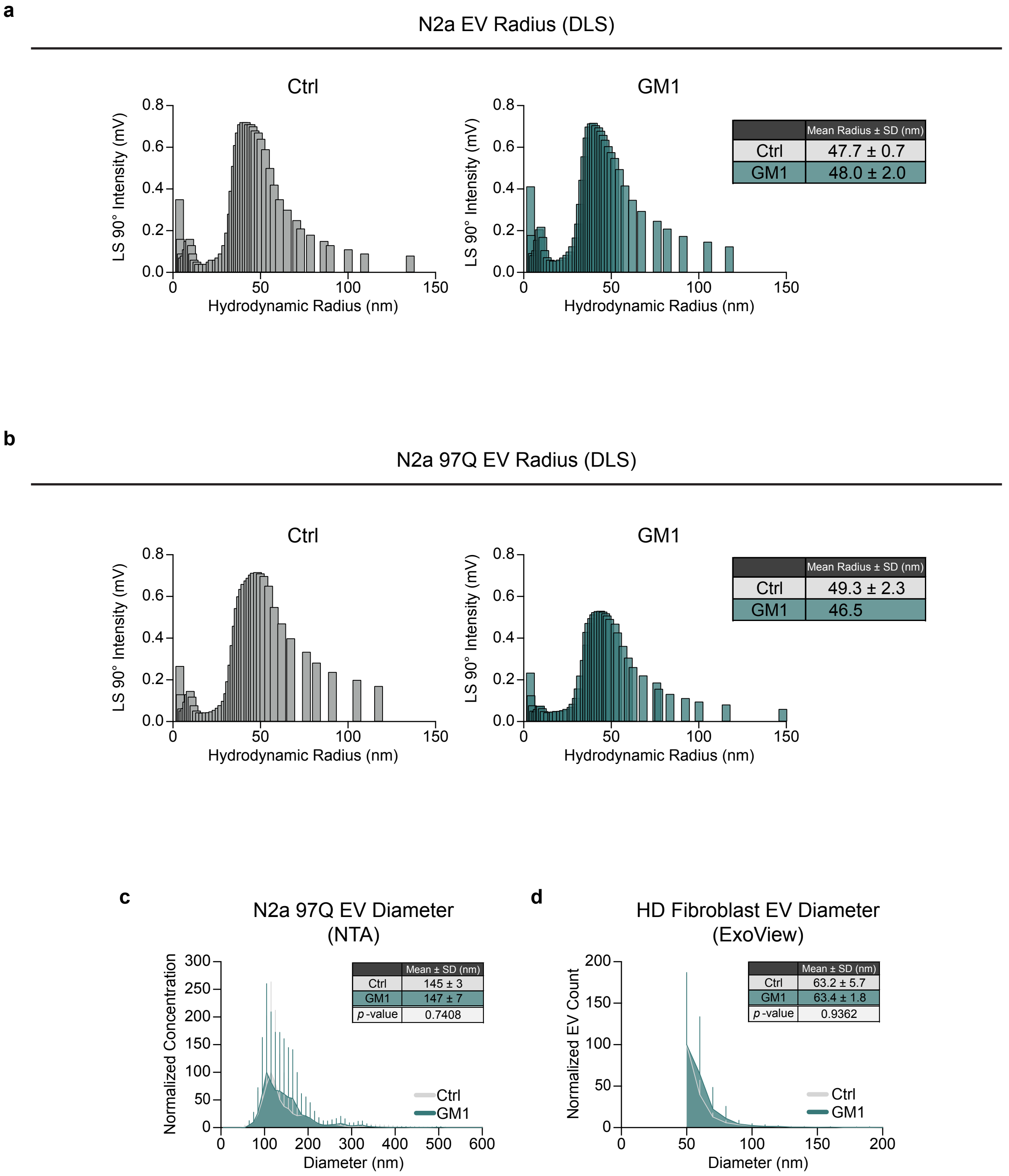

Supplementary figure 3

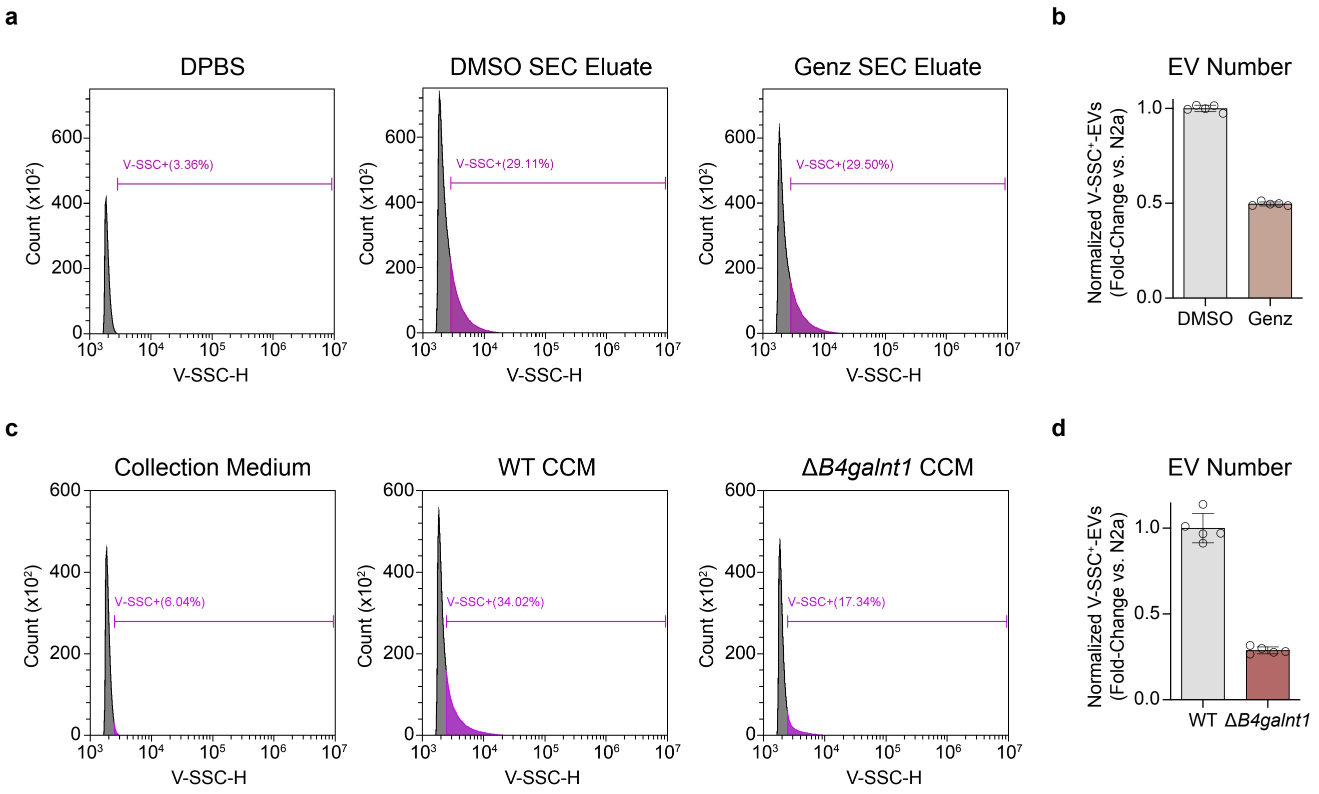

Supplementary figure 4

**a**

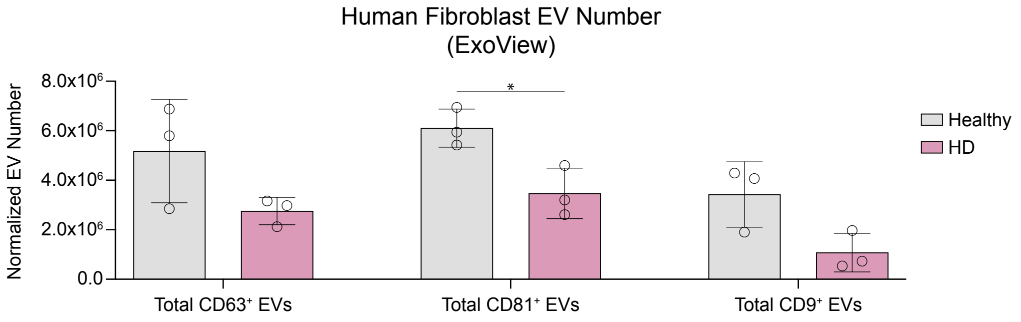


**b**

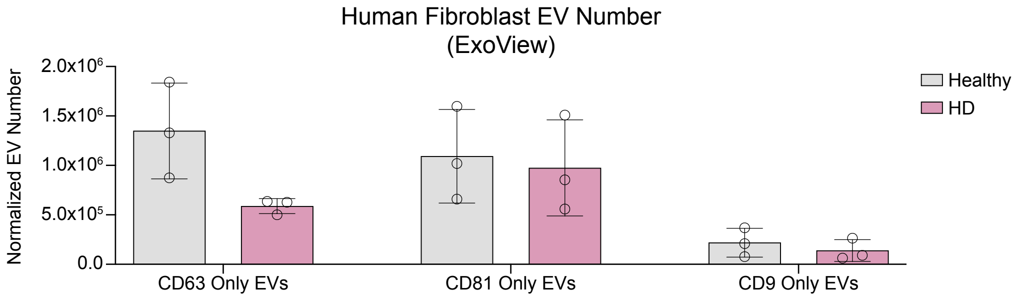


**c**

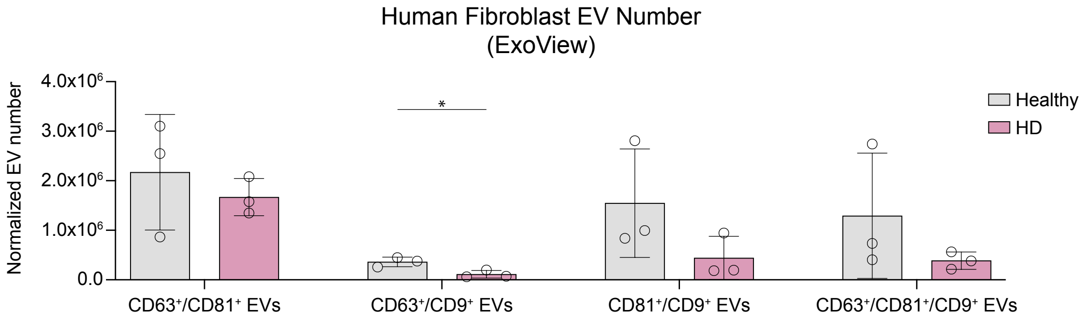

Supplementary figure 5

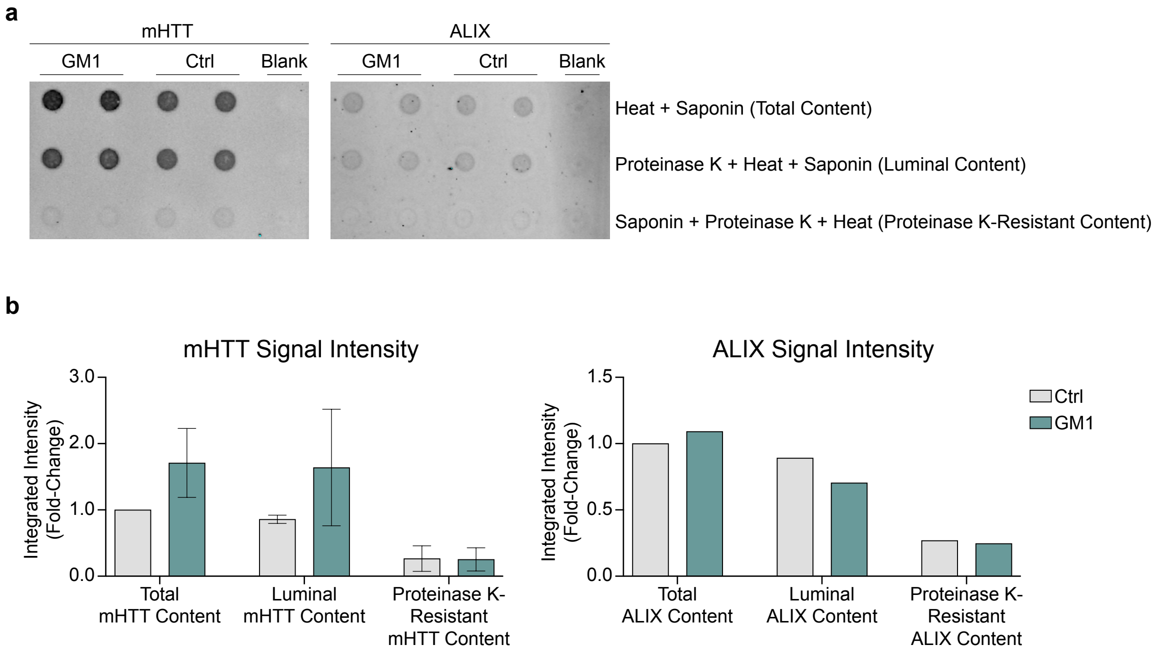
